# Activity of Estrogen Receptor β Agonists in Therapy-Resistant Estrogen Receptor-Positive Breast Cancer

**DOI:** 10.1101/2022.01.14.476328

**Authors:** Jharna Datta, Natalie Willingham, Jasmine M. Manouchehri, Patrick Schnell, Mirisha Sheth, Joel J. David, Mahmoud Kassem, Tyler A. Wilson, Hanna S. Radomska, Christopher C. Coss, Chad E. Bennett, Ramesh K. Ganju, Sagar D. Sardesai, Maryam Lustberg, Bhuvaneswari Ramaswamy, Daniel G. Stover, Mathew A. Cherian

**Author notes:** **Correspondence:** Mathew Cherian.

## Abstract

**Background:** Among women, breast cancer is the leading cause of cancer-related death worldwide. Estrogen receptor α positive (ERα+) breast cancer accounts for 70% of all breast cancer subtypes. Although ERα+ breast cancer initially responds to estrogen deprivation or blockade, resistance emergence compelling the use of more aggressive therapies. While ERα is a driver in ERα+ breast cancer, ERβ plays an inhibitory role in several different cancer types. To date, the lack of highly selective ERβ agonists without ERα activity has limited the exploration of ERβ activation as a strategy for ERα+ breast cancer.

**Methods:** We measured expression levels of ESR1 and ESR2 genes in immortalized mammary epithelial cells and different breast cancer cell lines. The viability of ERα+ breast cancer cell lines upon treatments with specific ERβ agonists, including OSU-ERb-12 and LY500307 was assessed. The specificity of the ERβ agonists, OSU-ERb-12 and LY500307, was confirmed by reporter assays. The effects of the agonists on cell proliferation, cell cycle, apoptosis, colony formation, cell migration, and expression of tumor suppressor proteins were analyzed. The expression of ESR2 and genes containing ERE-AP1 composite response elements was examined in ERα+ human breast cancer samples to determine the correlation between ESR2 expression and overall survival and that of putative ESR2 regulated genes.

**Results:** In this study, we demonstrate the efficacy of highly selective ERβ agonists in ERα+ breast cancer cell lines and drug-resistant derivatives. ERβ agonists blocked cell proliferation, migration and colony formation; and induced apoptosis and S and/or G2/M cell cycle arrest of ERα+ breast cancer cell lines. Also, increases in the expression of the key tumor suppressors FOXO1 and FOXO3a were noted. Importantly, the strong synergy between ERβ agonists and ERα antagonists suggested that the efficacy of ERβ agonists is maximized by combination with ERα blockade. Lastly, ESR2 (ERβ gene) expression was negatively correlated with ESR1 (ERα gene) and CCND1 RNA expression in human metastatic ER+/HER2-breast cancer samples.

**Conclusion:** Our results demonstrate that highly selective ERβ agonists attenuate the viability of ERα+ breast cancer cell lines in vitro and suggest that this therapeutic strategy merits further evaluation for ERα+ breast cancer.

## Introduction

Breast cancer is the most prevalent cancer among women globally (1). It is the second leading cause of cancer-related deaths among women. In 2020, there were 2.3 million new breast cancer cases and 685,000 breast cancer deaths worldwide. Despite advances in diagnostic procedures and improved therapies, globally breast cancer-related morbidity and mortality are on the rise. The majority of breast cancer-related deaths occur due to distant metastasis. About 60% of metastatic breast cancers (MBC) are estrogen receptor α positive (ERα+) and human epidermal growth factor receptor 2 non-amplified (HER2-) (2). Although the development of effective estrogen blocking agents and cyclin-dependent kinase 4/6 inhibitors (CDK4/6i) has doubled progression-free survival on first-line therapy of ERα+-HER2-MBC, endocrine and CDK4/6i resistance emerges causing disease progression. Appropriate post-CDK4/6i therapy is poorly defined due to incomplete understanding of CDK4/6i resistance, lack of effective agents and lack of clinical trials that address this important issue.

While augmented signaling through receptor tyrosine kinases, *NF1* loss, *C-MYC* amplification and activating mutations in the *ESR1* gene result in endocrine resistance, alterations of cell cycle genes cause CDK4/6i resistance (3-5). Due to redundancy and cross talk in these signaling pathways, attempts to counter therapeutic resistance by focusing on a single target have been mostly ineffective. Thus, there is an urgent need to develop novel therapeutic options in the second-line setting to improve the survival and response rate for this aggressive endocrine and CDK4/6i resistant MBC.

Estrogens play a vital role in breast tumorigenesis (6, 7). The stimulatory or repressive effects of estrogens are mediated through ERα and ERβ, which are gene products of *ESR1* and *ESR2*, respectively, and the G protein-coupled estrogen receptor (GPCR30). Unlike ERα, which has a clear oncogenic role in ERα+ breast cancer, ERβ behaves like a tumor suppressor in many biological contexts. For example, the tumor-suppressive function of ERβ was demonstrated through its knockdown in ERα+ cell lines, which induced an invasive phenotype, increased anchorage-dependent cell proliferation, and elevated EGF-R signaling (8). In the presence of estradiol, ERβ overexpression reduced cell proliferation *in vitro* and tumor formation *in vivo*, effects that are in contradistinction to those of ERα (9, 10). In these experiments ERβ also was shown to repress the expression of oncogenes such as c-myc and cyclin D1.

The transcriptional function of ERs involves their binding to estrogen response elements (ERE) within promoters and enhancers (11). There are multiple conformations of EREs in the human genome, including consensus and non-consensus EREs, single and multiple binding site, and composite EREs consisting of ERE half-sites in combination with binding sites for other transcription factors such as AP1 and Sp1. Although both the receptors exhibit transcriptional activity, they differ in their modes of transcriptional activation (12). Studies demonstrated that on certain E2 responsive ERE-AP-1 composite promoters, ERβ actually antagonizes the effects of ERα (13). For example, the cyclin D1 (*CCND1*) promoter, containing cAMP response element and an AP-1 binding site, is activated by estradiol in cells overexpressing ERα but is inhibited in cells overexpressing ERβ (13).

*ESR2* was discovered more than twenty years ago (14) but its clinical application was limited by the lack of highly selective ERβ agonists. Although, both ERα and ERβ are activated by binding to endogenous estrogens, the development of several highly selective synthetic ligands of ERα or ERβ has uncovered new avenues to probe the function of these receptors (15).

In the present study, we investigated the effects of a novel and highly selective ERβ selective agonist, OSU-ERb-12 (16), to inhibit preclinical models of ERα+ breast cancer and to counter endocrine and CDK4/6i resistance *in vitro*. We found that treatment of ERα+ breast cancer cell lines with OSU-ERb-12 caused apoptosis, induced cell cycle arrest (at S phase), as well as decreased cell proliferation, colony formation, and cell migration. FOXO1 and FOXO3a protein expression was significantly increased in cells treated with OSU-ERb-12, a potential mechanism for its tumor-suppressive effects (17).

## Materials and Methods

### Chemicals, drugs, plasmids, antibodies, primers and synthesis of MCSR-18-006

OSU-ERb-12 was synthesized in the Drug Development Institute (DDI) at OSU according to the procedure outlined before (16). LY500307 was also obtained from DDI, OSU. AC186 (cat# 5053), WAY200076 (cat# 3366), diarylpropionitrile (DPN; cat# 1494), 4-hydroxy-tamoxifen (Tam; cat# 3412/10), fulvestrant (Fas; cat# 10-471-0), and 1,3-*Bis*(4-hydroxyphenyl)-4-methyl-5-[4-(2-piperidinylethoxy)phenol]-1*H*-pyrazoledihydro-chloride (MPP; cat# 1991) were purchased from Tocris Bioscience. Elacestrant (RAD1901; cat# S9629) was purchased from *Selleckchem*. Abemaciclib (LY2835219; cat# 17740) was obtained from Cayman Chemical. Stock solutions (10 mmol/L) of the inhibitors were prepared in DMSO. CellTiter-Glo reagent (cat# G7570) and Dual-Luciferase Assay reagent (cat# E1960) were purchased from Promega Corporation. Lipofectamine 3000 was obtained from Thermo Fisher Scientific.

pRLTK plasmid was obtained from Promega. 3XERE TATA luc (luciferase reporter that contained three copies of vitellogenin Estrogen Response Element) was a kind gift from Donald McDonnell (Addgene plasmid # 11354; http://n2t.net/addgene: 11354; RRID: Addgene_11354). Plasmids expressing pcDNA3 (OHu23619C; pcDNA3.1+: RRID: Addgene_10842), ERβ (OHu25562C; pcDNA3.1+), c-Flag pcDNA3 (OHu23619D; pcDNA3.1+/C-(K) DYK), c-Flag ERα (OHu26586D; pcDNA3.1+/C-(K) DYK), and c-Flag ERβ (OHu25562D; pcDNA3.1+/C-(K) DYK were obtained from GenScript.

Antibodies to ERα (D8H8; 8644), FOXO1 (D7C1H; cat# 14952, RRID:AB_ 2722487), FOXO3a (75D8; cat# 2497), PARP (cat# 9542, RRID:AB_2160739), cleaved PARP (Asp24, D64E10; cat# 5625, RRID:AB_10699459), caspase-3 (8G10; cat# 9665, RRID:AB_2069872), cleaved caspase-3 (D175; cat# 9664, RRID:AB_2070042), caspase-7 (cat# 9492, RRID:AB_2228313), cleaved caspase-7 (asp198, D6H1; cat# 8433, RRID:AB_11178377), and GAPDH (D16H11; cat# 8884, RRID:AB_11129865) were obtained from Cell Signaling Technology. Antibodies against ERβ (clone 68-4; cat# 05-824) and M2 Flag (cat# F1804) were purchased from Sigma-Aldrich. The following primers were used for the corresponding mRNAS.

ESR2 full length: forward (5’- CTCCAGATCTTGTTCTGGACAGGGAT-3’), reverse (5’- GTTGAGCAGATGTTCCATGCCCTTGTTA-3’); ESR2 all isoforms: forward (5’-ACACA CCTTACCTGTAAACAGAGAG-3’), reverse (5’- GGGAGCCACACTTCACCATTCC-3’); ESR1: forward (5’-CCGCCGGCATTCTACAGGCC-3’), reverse (5’-GAAGAAGGCCTTG CAGCCCT-3’); GAPDH: forward (5’-GTCGTATTGGGCGCCTGGTC-3’), reverse (5’-TT TGGAGGGATCTCGCTCCT-3’).

^1^H-NMR spectra were recorded using a Bruker AV300NMR, AVIII400HD NMR spectrometer or a DRX400 NMR spectrometer at The Ohio State University College of Pharmacy. Chemical shifts (*δ*) are specified in ppm from chemical reference shifts for internal deuterated chloroform (CDCl_3_) set to 7.26 ppm. Coupling constants are defined in Hz. Mass spectra were obtained using an Advion Expression Model S Compact Mass Spectrometer equipped with an APCI source and TLC plate express or using a Thermo LTQ Orbitrap mass spectrometer. For carborane-containing compounds, the obtained mass resembling the most intense peak of the theoretical isotopic pattern was described. Measured patterns corresponded with calculated patterns. Unless otherwise noted, all reactions were carried out under argon atmosphere using commercially available reagents and solvents. Details of the procedure for the synthesis of MCSR-18-006 has been provided in Supplemental Data.

### Cell culture, cell viability and generation of resistance

Normal mammary epithelial cells MCF10A (ATCC Cat# CRL-10317, RRID:CVCL_0598), breast cancer cell lines, MCF7 (ATCC HTB-22), T47D (ATCC HTB-133; : NCI-DTP Cat# T-47D, RRID:CVCL_0553), ZR-75-1 (ATCC CRL-1500), MDA-MB 231 (ATCC HTB-26, RRID:CVCL_0062), MDA-MB 468 (ATCC HTB-132, RRID:CVCL_0419) and HEK-293T (ATCC Cat# CRL-3216, RRID:CVCL_0063) were obtained from ATCC. All the cells were grown according to supplier’s recommendation in a humidified atmosphere containing 5% CO_2_ at 37^0^C. Cells were passaged and media changed every 2 days. Mycoplasma contamination of the cells were checked monthly using the MycoAlert Plus Mycoplasma Detection Kit (cat# LT07-703) (Lonza) following the manufacturer’s protocol. For routine experiments, parental and drug resistant cells of MCF7 and T47D were cultured in phenol red-free basal medium (DMEM) media, containing charcoal-stripped fetal bovine serum along with other additives as recommended.

T47D cells were treated gradually at increasing concentrations with 4-hydroxy-tamoxifen (Tam), fulvestrant/Faslodex (Fas; estrogen receptor antagonist) or abemaciclib (cyclin dependent kinase 4/6 inhibitor; CDK4/6i) to generate resistant cell lines (T47D-TamR, T47D-FasR and T47D-CDK4/6iR). Similarly, MCF7 cells were treated with increasing concentrations of abemaciclib to generate MCF7-CDK4/6iR cells. Control cells were treated with the vehicle DMSO. To evaluate the development of resistance, cells were examined for viability every 4 to 6 weeks with the CellTiter-Glo assay (Promega). Cell viability was measured in quadruplicates by seeding the cells (2,000 to 3,000 per well in 96-well plate), followed by addition of Tam, Fas, or abemaciclib at different dilutions or DMSO (vehicle control) after 24 hours. Seventy-two hours later, luminescence was measured after addition of CellTiter-Glo reagent following the manufacturer’s protocol. Cell viability was calculated as percentage relative to vehicle controls (100%). Viability curves were plotted using GraphPad Prism software (GraphPad Prism, RRID: SCR_002798). Upon manifesting resistance, cells were maintained with continued drug exposure at concentrations to which they were resistant.

Immortalized mammary epithelial MCF10A cells as well as MCF7 and T47D breast cancer (parental and respective resistant) cells were plated (in quadruplicates) in 96-well plates (2000-3000 cells/well) and allowed to grow overnight followed by treatment with OSU-ERb-12, LY500307, DPN (Diarylpropionitrile), AC186, WAY200070 (WAY) at varying concentrations as indicated. The fresh medium and drugs were replaced every alternate day. Cell viability was assessed after 7 days of initial drug exposure using CellTiter-Glo Luminescent Cell Viability Assay and the viability curves were plotted as mentioned above.

### *Reverse Transcription Polymerase Chain Reaction (RT-PCR), western blot analysis, and* Estrogen Response Element Luciferase (ERE-LUC) reporter assays

Total RNA was isolated from cells using TRIzol reagent (cat# 15596026) (Invitrogen, CA) following the manufacturer’s instructions, treated with DNase 1 and reverse transcribed into cDNA using high capacity cDNA reverse transcription kit (Applied Biosystems, Foster City, CA). Real-time RT-PCR (qRT-PCR) was performed using 0.01-0.05μg cDNA with SYBR Green mastermix (Applied Biosystems) in an Applied Biosystems thermocycler. The fold difference in target gene mRNA levels normalized to GAPDH was calculated using the ΔΔCT method. Semi-quantitative PCR was performed using the same set of primers as in qRT-PCR and visualized after electrophoretic separation to confirm the identity of the amplicons. The primers were designed spanning exon-exon junction to avoid non-specific amplification of genes.

Whole cell extracts were prepared in cell lysis buffer (50 mM Tris pH 8.1, 10 mM EDTA, 1% SDS, and 1% IGEPAL (CA-630, 18896; Sigma–Aldrich) followed by sonication and centrifugation at 14,000 rpm for 10 mins at 4°C. Protein concentrations in the extracts were measured using the bicinchoninic acid (BCA) method using BSA as the standard. Equivalent amounts of protein from whole cell lysates were mixed with 4× Laemmli’s buffer, boiled for 5 minutes at 97°C, separated by SDS-polyacrylamide (10%) gel electrophoresis (Thermo Fisher Scientific), transferred to nitrocellulose membranes (GE Healthcare, Chicago, IL) and probed with the antibodies described above. Membranes were incubated overnight at 4°C with the primary antibody, washed and blotted for an hour with secondary anti-mouse/rabbit (HRP-conjugated) antibodies). Enhanced chemiluminescence substrate detection system (Millipore-Sigma) was applied to detect bound antibody complexes and visualized by autoradiography. The loading control was GAPDH. The intensity of the protein bands was quantified using image studio (Licor). HEK293T cells (7.5 × 10^4^/well) seeded in a 24-well plate were transfected for 12 hours with ERE-Luc, pRLTK (internal control, Promega), and c-Flag pcDNA3/ERα/ERβ plasmids using Lipofectamine 3000 (Thermo Fisher Scientific) following manufacturer’s protocol. The media was changed with phenol-red free DMEM containing 10% charcoal-stripped FBS, and insulin (6ng/mL). Six hour later cells were treated with OSU-ERb-12, or LY500307 at varying concentrations as indicated. DMSO was used as a vehicle control. Luciferase activity was assessed after 72 hours of transfection using Dual-Luciferase Assay System (Promega).

### Cell proliferation, cell cycle analysis, apoptosis, clonogenic survival, and cell migration assays

MCF7 and T47D cells were plated at 5×10^5^ cells per plate in phenol red-free complete DMEM supplemented with charcoal-stripped FBS. The cells were treated for 72 hours with OSU-ERb-12(0.5 μmol/L and 10 μmol/L) or LY500307 (MCF7: 0.5 μmol/L and 3 μmol/L; T47D: 0.5 μmol/L and 7 μmol/L). Differing concentrations were used to avoid complete loss of viability. DMSO and fulvestrant (0.5 μmol/L) were used as negative and positive controls, respectively. The cells were harvested and stained as per the protocol for the Click-iT Edu Alexa Fluor 647 kit (Invitrogen; cat# C10424). The stained cells were analyzed via flow cytometry (BD FACSCalibur Flow Cytometer).

MCF7 and T47D cells were plated in 100 mm dishes (5×10^5^ cells) in phenol red-free complete DMEM supplemented with charcoal-stripped FBS. The cells were treated for 72h hours with OSU-ERb-12 or LY500307 at the indicated concentrations. DMSO was used as vehicle control. The cells were harvested, fixed in 70% ethanol and stained with propidium iodide. The stained cells were analyzed via flow cytometry on a BD FACSCalibur Flow Cytometer.

Breast cancer MCF-7 and T47D cells were plated and treated 24h later with OSU-ERb-12 (0.5 μmol/L and 10 μmol/L) or LY500307 (MCF7: 0.5 μmol/L and 3 μmol/L; T47D: 0.5 μmol/L and 7 μmol/L) for 48 hours. Cells were collected and processed according to the manufacturer (TUNEL Assay Kit - BrdU-Red (cat# ab66110) (Abcam). Processed breast cancer cells were analyzed on BD FACSCalibur Flow Cytometer.

MCF7 and T47D cells were plated in 6-well plates (1∼2×10^4^ cells/well). Twenty four hours after plating, cells were treated with OSU-ERb-12, LY500307, or vehicle (DMSO) for 7-10 days. The fresh medium and drugs were replaced every other day. Next, cell colonies were washed with PBS, fixed with paraformaldehyde (4%), and stained with crystal violet solution (0.05%). Colonies were then washed with water and air-dried. Visible colonies were counted manually.

MCF7 Cells were seeded, treated with DMSO (control), OSU-ERb-12 or LY500307 and allowed to grow until confluence. Confluent monolayers were scratched using a sterile pipette tip, washed and incubated in complete medium containing DMSO or the drugs. Plates with similar scratch were selected by examination under microscope and used for further analysis. Images were captured immediately after scratch (0 hour) and 24 hours post-scratch. Migration of cells from the edge of the groove toward the center was monitored at 24 hours (40 magnification). To calculate the fraction of the gap covered by the cells in a 24-hour period the width of the scratch was measured at 0 hour and at 24 hours. Mean fraction of filled area was determined and data presented was normalized to the control cells.

### Messenger RNA expression of patient samples and Statistical and bioinformatics analyses

Patients treated at The Ohio State University Comprehensive Cancer Center – Arthur G. James Cancer Hospital and Richard J. Solove Research Institute since 1998 with a diagnosis of metastatic ERα+ and HER2 negative (ERα+/HER2^-^) breast cancer and confirmed RNA sequencing analysis were eligible for this retrospective clinical correlation. Following IRB approval (OSU 1999C0245), the list of patients fulfilling the previous criteria was obtained from the Ohio State University Medical Center and James Cancer Registry. 118 medical record were reviewed and 37 patients had RNA sequencing performed through the Oncology Research Information Exchange Network (ORIEN) and were deemed eligible.

Data for the 37 eligible patients were initially queried and obtained from The Ohio State University Information Warehouse and from ORIEN-AVATAR and uploaded into REDCap (REDCap, RRID:SCR_003445). Data missing from the initial query were populated using manual review of each patient’s electronic medical record.

The objective was to determine the mRNA expression levels of the genes which are targets of ER-AP1 mediated transcription and AP1 independent ER mediated transcription including CCND1, MYC, IGF-1, Bcl-2, MMP-1, FN1; IGFBP-4, E2F4, CXCL12, PGR, EBAG9, and TRIM25 and correlated with ERα and ERβ.

Viability, proliferation, apoptosis, and cellular mRNA expression were analyzed using students t-test.

For each dose, linear mixed models were fit for log-transformed viability with fixed effects for regimen (4-hydroxy-tamoxifen, OSU-ERb-12 and 4-hydroxy-tamoxifen+OSU-ERb-12) and random effects accounting for within-batch correlation of replicates. Predictions and standard errors for viability of the 4-hydroxy-tamoxifen+OSU-ERb-12 combination under a hypothesized Bliss independence model were computed from estimated mean viabilities under 4-hydroxy-tamoxifen and OSU-ERb-12 alone via the formula Log_Viability (Bliss) = Log_Viability(4-hydroxy-tamoxifen) + Log_Viability (OSU-ERb-12). Interaction at each dose was quantified as the ratio of the predicted viability under the Bliss independence model over the estimated viability under the tested 4-hydroxy-tamoxifen + OSU-ERb-12 combination, with ratios >1 indicating synergy.

Total RNA was sequenced with minimum 20 million reads and >65% reads aligned identified for subsequent processing to transcript abundance values (FKPM; fragments per kilobase per million reads) following ORIEN standard pipeline: STAR aligner (STAR, RRID:SCR_004463), Star-fusion, and RSEM (RSEM, RRID:SCR_013027) with genome GRCh38 alignment/annotation. Statistical analysis was performed using the R statistical software, including the ‘survival’ package. Summary statistics were computed for demographic variables and expression levels (FPKM), and Spearman correlation coefficients were computed for *ESR1* (ERα) and *ESR2* (ERβ) versus other expression levels. Cox regression was used to calculate the association between overall survival and log2(1 + FPKM) for ERα and ERβ expression levels.

## Results

### Selection for drug-resistant MCF7 and T47D cell lines

We cultured MCF7 and T47D cell lines, in the presence of DMSO (control), 4-hydroxy-tamoxifen, fulvestrant, or the CDK4/6i abemaciclib at gradually increasing concentrations to select for acquired resistance. With extended exposure, both the cell lines demonstrated decreased sensitivity to the drugs compared with the corresponding parental controls **(Supplemental Fig. S1)**. Chemical Structures of the drugs/inhibitors used in this study have been provided in **Supplemental Fig. S2**.

Lack of activation of ERE-luciferase reporter vector by overexpressed ERα and ERβ proteins in 293T cells treated with the inactive chemical analog of OSU-ERb-12, MCSR-18-006, is shown in **Supplemental Fig. S3**. The lack of binding affinity of MCSR-18-006 for ERα and ERβ proteins as measured by radiolabeled estradiol competition binding assays is shown in **Supplemental Fig. S4**.

### ESR2 and ESR1 genes and their protein products are differentially expressed in ERα+ parental and resistant as well as triple-negative breast cancer cell lines, and ERβ driven ERE-luciferase promoter activity is significantly enhanced upon treatment with selective ERβ agonists compared to that of ERα

We assessed the basal expression levels of *ESR2* and *ESR1* in three ER-positive breast cancer cell lines (MCF7, T47D and ZR-75-1), the derivative endocrine-resistant and CDK4/6i resistant lines (of MCF7 and T47D) and compared with those of immortalized mammary epithelial cells (MCF10A) (**Fig.1**) using primers that selectively amplified only the full-length, canonical *ESR2*, or that amplified all known splice variants of *ESR2* (**Supplemental Fig. S5A)**, as well as primers that specifically amplify full-length *ESR1*. The p-value**s** and 95% confidence interval (CI) of corresponding expression data are shown in **Supplemental Table 1**. qRT-PCR data demonstrated a comparable expression of full-length *ESR2* in MCF7 and MCF10A lines (**Fig. 1A, Supplemental Table 1)**. While MCF7-FasR and MCF7-CDK6-O/E cells displayed no significant increase in full length *ESR2* expression relative to the control MCF10A cells, MCF7-TamR, and MCF7-CDK4/6iR cells showed 3.6-fold (p=0.0035) and 6-fold (p=0.0001) higher expression levels (**Fig. 1A, Supplemental Table 1**). On the other hand, T47D exhibited a 4.8-fold (p=0.0265) higher expression of *ESR2* compared to MCF10A cells. Significantly higher expression of full-length ESR2 in T47D-TamR (5.1-fold, p=0.0009) and T47D-CDK4/6iR (5.1-fold, p=0.0075) compared to MCF10A was noted (**Fig. 1A, Supplemental Table 1**). ZR-75-1 cells displayed the highest level of full-length *ESR2* RNA expression (∼19-fold higher than MCF10A; p< 0.01) **(Fig. 1A, Supplemental Table 1**). Both the TNBC lines had significantly higher expression of full-length *ESR2* compared with MCF10A (MDA-MB-231: 4.4-fold, p<0.05; MDA-MB-468: 5.2-fold, p<0.01) and these levels were comparable to those in the ERα+ MCF7 and T47D breast cancer cell lines.

**Figure 1:**
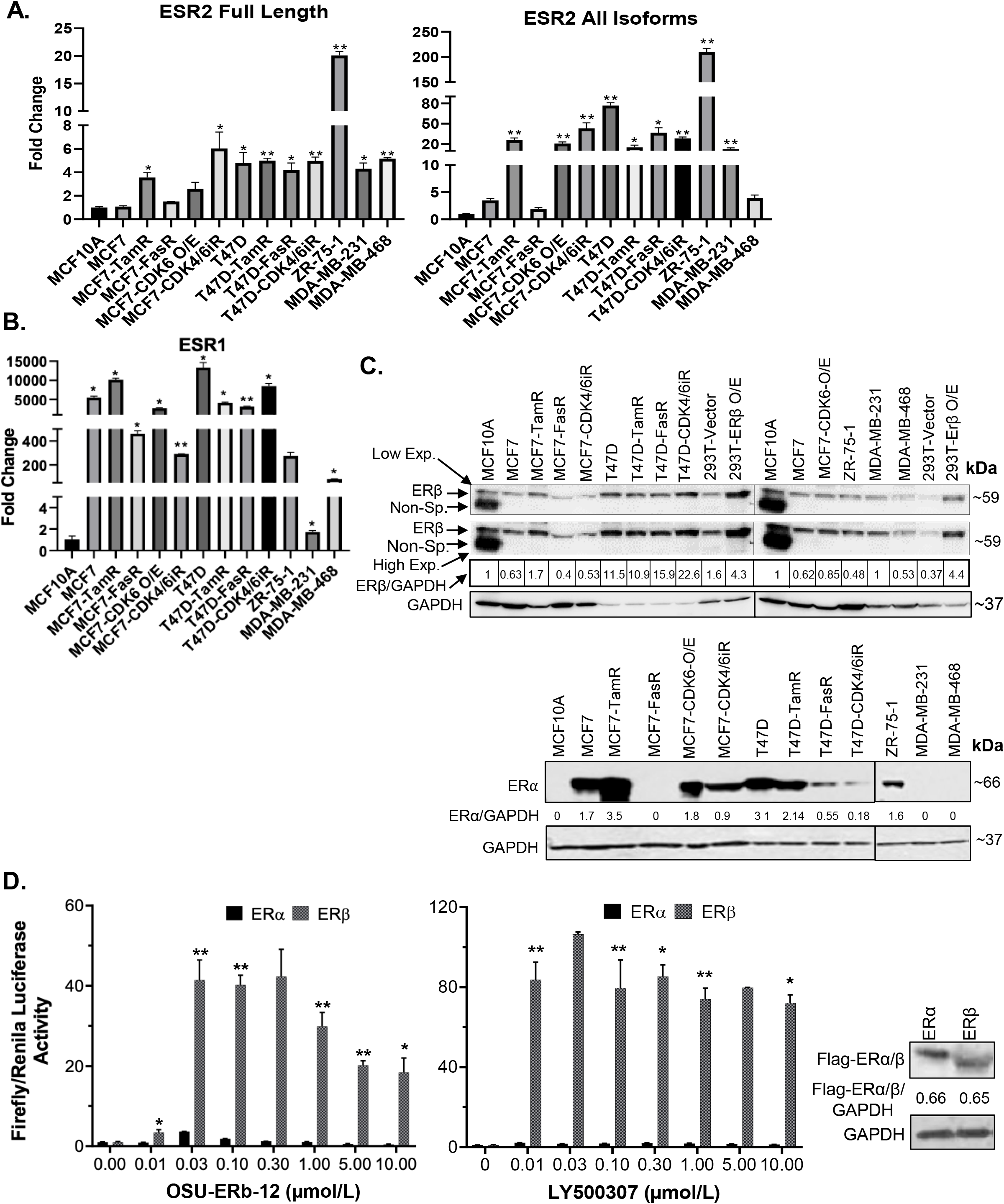
A-C, ESR1 and ESR2 genes are differentially expressed in ERα+ parental, respective endocrine resistant, and triple negative breast cancer cell lines. **A** and **B**, Expression of *ESR1* and *ESR2* in immortalized mammary MCF10A, transformed ERα+ MCF7 and T47D, endocrine resistant MCF7-TamR, MCF7-FasR, T47D-TamR, and T47D-FasR, CDK6 over-expressing MCF7 (MCF7-CDK6 O/E), CDK4/6 inhibitor resistant MCF7 (MCF7-CDK4/6iR) and T47D (T47D-CDK4/6iR), ZR-75-1, and triple negative breast cancer (TNBC; MDA-MB231, MDA-MB-468, Hs578t) cell lines. Total RNA was isolated from the established cell lines using TRIzol. The expression of each gene was assessed by quantitative RT-PCR (qRT-PCR) performed with the DNase-treated RNA samples using gene-specific primers spanning exon-exon junctions that include large introns in the corresponding genomic sequence to avoid genomic DNA amplification. Gene expression was calculated by ΔΔCt method using GAPDH as an internal control. The expression of each gene is shown as the fold change relative to MCF10A. All reactions were done in triplicate and the experiment was repeated twice. Data were plotted as mean ± SD. **A**, *ESR2* genes; full length (left) and all isoforms (right). **B**, *ESR1*. **C**, whole-cell lysates were extracted and immunoblot analyses were performed for ERβ and GAPDH (loading control) (upper panel), and ERα and GAPDH (lower panel). Intensity of the protein bands was quantified using Image Studio (LiCor) software. Numbers under the lanes of each cell line represent normalized values of the corresponding protein band (ERβ or ERα). Normalized band intensity of MCF10A was considered as 1. Immunoblot analyses were repeated twice with corresponding biological replicates. Reproducible results were obtained in each independent experiment. GAPDH, glyceraldehyde-3-phosphate dehydrogenase. For ERβ (upper panel) two different exposures were provided; low exp.= low exposure; high exp.= higher exposure of the blot **D, ERE-Luciferase driven promoter activity upon treatment with selective ERβ agonists is significantly higher in ectopically expressing cells with ERβ compared to that of ERα**. HEK293T cells were transfected with c-Flag pcDNA3 (vector control), c-Flag ERα or c-Flag ERβ in combination with ERE-Luciferase (reporter) and TK-renilla (pRLTK; internal control) plasmids (as described in Materials and Methods section). Forty eight hours after treatment of the cells with ERβ specific agonists Renilla and Firefly luciferase activities were measured using the dual-luciferase reporter assay system. Renilla luciferase was normalized to Firefly luciferase. Treatment with: OSU-ERb-12 (0-10 μmol/L) (left) and LY500307 (0-10 μmol/L) (middle). Each assay was performed in triplicate with three experimental replicates. (mean ±SD, *: p<0.05, **: p<0.01). Right panel shows equal expression of ERα and ERβ as determined by western blot analysis using anti-flag antibody. Intensity of Flag-ERα/ERβ was normalized to GAPDH. The numbers under the corresponding protein band represent normalized values of the corresponding protein band intensity.

When we measured expression levels using primers that amplified all the splice isoforms of *ESR2*, the expression levels were significantly higher than MCF10A in most of the cells tested except MCF7, MCF7-FasR, and the TNBC line MDA-MB-468 (**Fig. 1A, Supplemental Table 1)**. About 5,000 (p<0.05) and 12,000-fold (p<0.05) increased *ESR1* expression was noted in MCF7 and T47D cells, respectively, compared to MCF10A (**Fig. 1B, Supplemental Table 1**).

To check the specificity of the primers to amplify the correct PCR products we performed agarose gel electrophoresis with the samples of qRT-PCR. Our data showed a single band (**Supplemental Fig. S5B**) with correct PCR products that were confirmed by sequencing.

Next, we performed western blot analyses to evaluate the expression of full-length ERβ and ERα proteins with the cell lysates (**Fig. 1C**). We tested antibodies raised against ERβ from several different sources including Developmental Studies Hybridoma Bank (CWK-F12), Invitrogen (PPZ0506), and Sigma (clone 68-4). Of these tested antibodies while CWK-F12 and PPZ0506 were specific but only sensitive to the overexpressed (positive control) ERβ protein, the antibody from Sigma was specific as well as sensitive to ERβ protein expressed at endogenous levels. As shown in **Fig. 1C (upper panel)**, all the parental and resistant ERα+ cell lines, TNBC lines as well as immortalized mammary epithelial cells expressed full-length ERβ. As expected, our data demonstrated that all the ERα+ parental cell lines but none of the TNBC cell lines expressed ERα protein. MCF7-TamR cells expressed more ERα protein than the parental MCF7 cells while MCF7-FasR had no detectable ERα expression. Similarly, T47D-FasR and T47D-CDK4/6iR cells had lower expression of ERα than the parental T47D cells.

In summary, full-length ERβ mRNA and protein is expressed in ERα+ breast cancer cell lines at levels that are comparable to expression levels in TNBC cell lines, and its expression is preserved in all the resistant derivative cell lines.

To determine the specificity of ERβ agonists, we treated HEK293T cells with OSU-ERb-12 or LY500307 (known selective ERβ agonist) at increasing concentrations following co-transfection with plasmids 3XERE TATA luc, pRLTK, FLAG-ERα or FLAG-ERβ (please see Materials and Methods section for details), and measured luciferase reporter activity (**Fig. 1D**). The expression of FLAG-ERα and FLAG-ERβ proteins was similar as measured by immunoblot for FLAG performed on lysates from the vehicle-treated 293T cells transfected with the corresponding expression plasmids (**Fig. 1D, right panel**). Comparison of the induction of luciferase activity demonstrated that ERα exhibited full activity in presence of 30 nmol/L OSU-ERb-12 and 10 nmol/L LY500307 treatment. Our data showed that luciferase activation by OSU-ERb-12 was significantly increased in the ERβ expressing cells as compared to those that expressed ERα. For example, at 30 nmol/L of OSU-ERb-12 there was ∼4-fold (p<0.05) and ∼40-fold (p<0.05) increase in luciferase activity, respectively, compared to their corresponding vehicle-treated cells (**Fig. 1D, left panel**). There was 10-fold (p=0.0059) higher ERE-LUC activity in ERβ overexpressing cells compared to that of ERα by OSU-ERb-12 at 30 nmol/L (**Supplemental Table 2**). Similarly, for LY500307 at 10 nmol/L there was 2.1–fold (maximum induction; p<0.05) activation by ERα and 84-fold (p<0.05) activation by ERβ compared to the corresponding vehicle-treated samples (**Fig. 1D, central panel, Supplemental Table 2**). At this concentration of LY500307, ERβ demonstrated 40-fold higher activity (p=0.0038) compared to ERα.

### Cell viability assay data demonstrates significant cytotoxicity of the selective ERβ agonists and those synergize with ERα agonists to demonstrate cytotoxicity towards ERα+ breast cancer cell lines

Next, we assessed the viability of parental, endocrine resistant, CDK4/6i-R MCF7 and T47D, and MCF7-CDK6 O/E cell lines following treatment with ERβ agonists OSU-ERb-12 and LY500307 (**Fig. 2, Supplemental Table 3**). We assessed Cell viability after 7 days of initial drug exposure using CellTiter-Glo Luminescent Cell Viability Assay. This duration is consistent with that used for toxicity assays with other endocrine agents such as fulvestrant (18, 19). We compared the viability of the drug treated transformed cell lines to that of MCF10A cells. The IC50 values for T47D cells (OSU-ERb-12: 10.43μmol/L-**Fig. 2C**; LY500307: 7.29 μmol/L-**Fig. 2D**), tamoxifen and fulvestrant resistant MCF7 cells, tamoxifen and fulvestrant resistant T47D cells, CDK6 overexpressing MCF7 cells, abemaciclib resistant MCF7 cells and abemaciclib resistant T47D cells were significantly lower than that of MCF10A cells (OSU-ERb-12: 13.96 μmol/L ; LY500307: 30.53 μmol/L; **Fig. 2, Supplemental Table 3)**. Compared to the parental MCF7 cell line, all the resistant lines except MCF7-CDK6 O/E had significantly lower IC50 values for OSU-ERb-12 (**Fig. 2A)**. Similarly, all three resistant T47D lines displayed significantly higher sensitivity towards OSU-ERb-12 compared to their parental counterpart (**Fig.2C, Supplemental Table 3)**.

**Figure 2:**
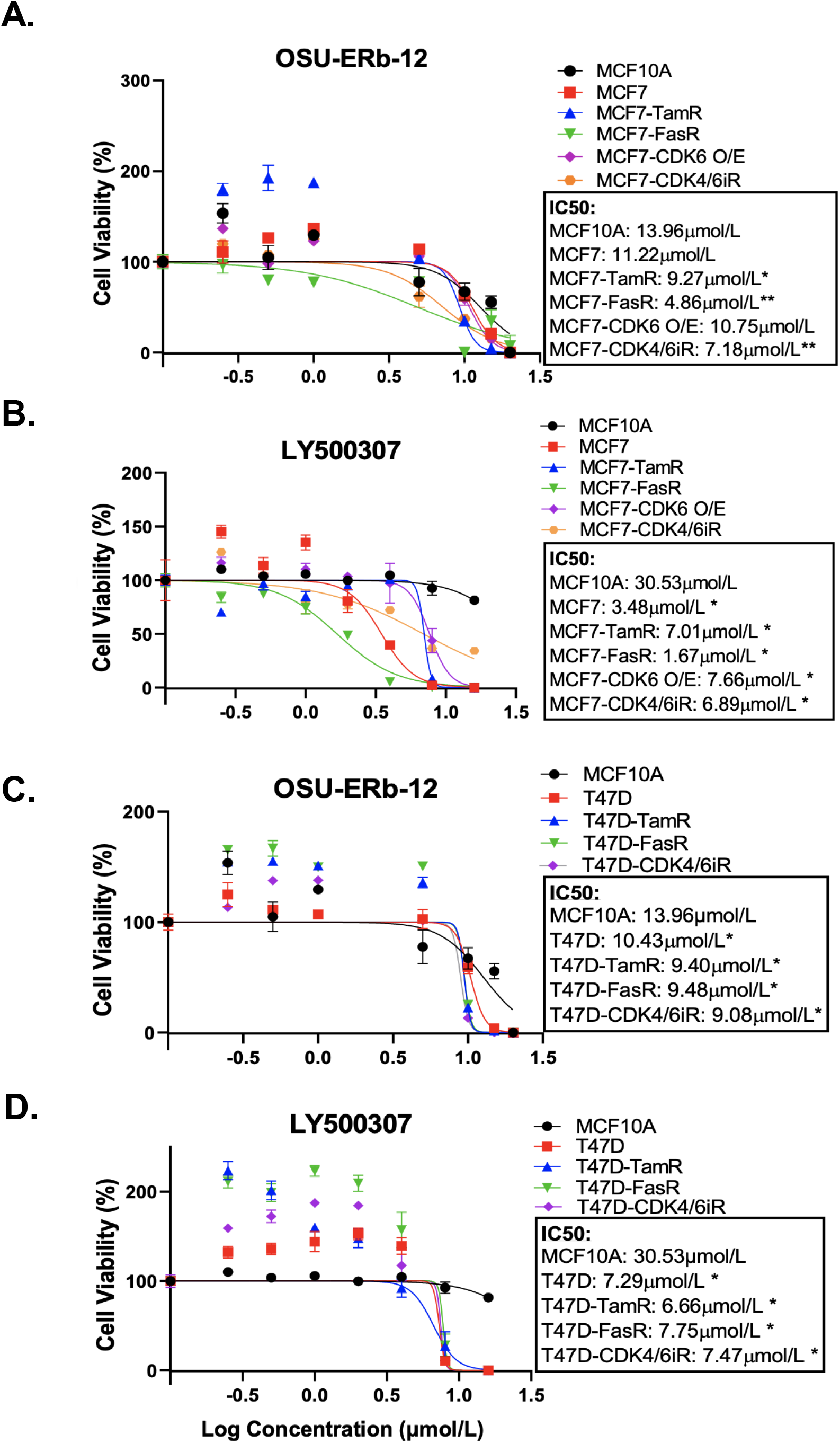
Selective ERβ agonists demonstrate significant cytotoxicity in ERα+ parental and respective endocrine resistant breast cancer cell lines. Cytotoxicity assays were performed on immortalized MCF10A, ER positive MCF7 and T47D, endocrine resistant MCF7 and T47D, CDK4/6 inhibitor resistant MCF7 and T47D, and CDK6 over-expressing MCF7 (MCF7-CDK6 O/E) cells. Viable cells were measured after seven days of treatment with DMSO (control) or the drugs at the indicated concentrations using CellTiterGlo assay. The percentage of viable cells is shown relative to DMSO vehicle-treated controls (mean ± SD, *: p<0.05, **: p<0.01). Assays were performed in quadruplicates (three experimental replicates). Cell viability assay performed after treatment with: **A & C**, OSU-ERb-12 **B & D**, LY500307. TamR= Tamoxifen resistant, FasR=Fulvestrant resistant, CDK6 O/E= CDK6 overexpressing, CDK4/6iR= CDK4/6 inhibitor resistant, MPP= methyl-piperidino-pyrazole.

Despite a high degree of selectivity, we saw some activation of ERα by both ERβ agonists in our reporter assay (**Fig. 1D**). We also observed an increase in viability of ERα+ breast cancer cell lines when exposed to low concentrations of both ERβ agonists. We hypothesized that combining ERβ agonists with an ERα antagonist would increase their activity and eliminate their stimulatory effects at low concentrations. We tested several ERα antagonists, namely, 4-hydroxy-tamoxifen (selective estrogen receptor modulator), fulvestrant, elacestrant (both selective estrogen receptor degraders/SERDs), and MPP (selective ERα antagonist) at concentrations that fully block ERα in combination with OSU-ERb-12. As shown in **Fig. 3A & 3B**, in T47D cells, all these ERα antagonists caused a significant reduction in the IC50 of OSU-ERb-12 and eliminated its stimulatory effects at low concentrations. Of the tested drugs, 4-hydroxy-tamoxifen, when used at a concentration of 0.5 μmol/L, displayed the highest efficacy leading to the reduction of IC50 for OSU-ERb-12 to 1 μmol/L from 14.10 μmol/L **(Fig. 3A, Supplemental Table 4**). We further analyzed the validity of the combination treatment of OSU-ERb-12 and 4-hydroxy-tamoxifen using the Bliss independence model (please see Materials and Methods for details). Our data demonstrated a significant dose-response with synergy (**Fig. 3C, Supplemental Table 4)**. There was evidence of synergy (the ratio being 1 or above) at all doses for the combination of OSU-ERb-12+Tam. There was no evidence of antagonism at any dose.

**Figure 3:**
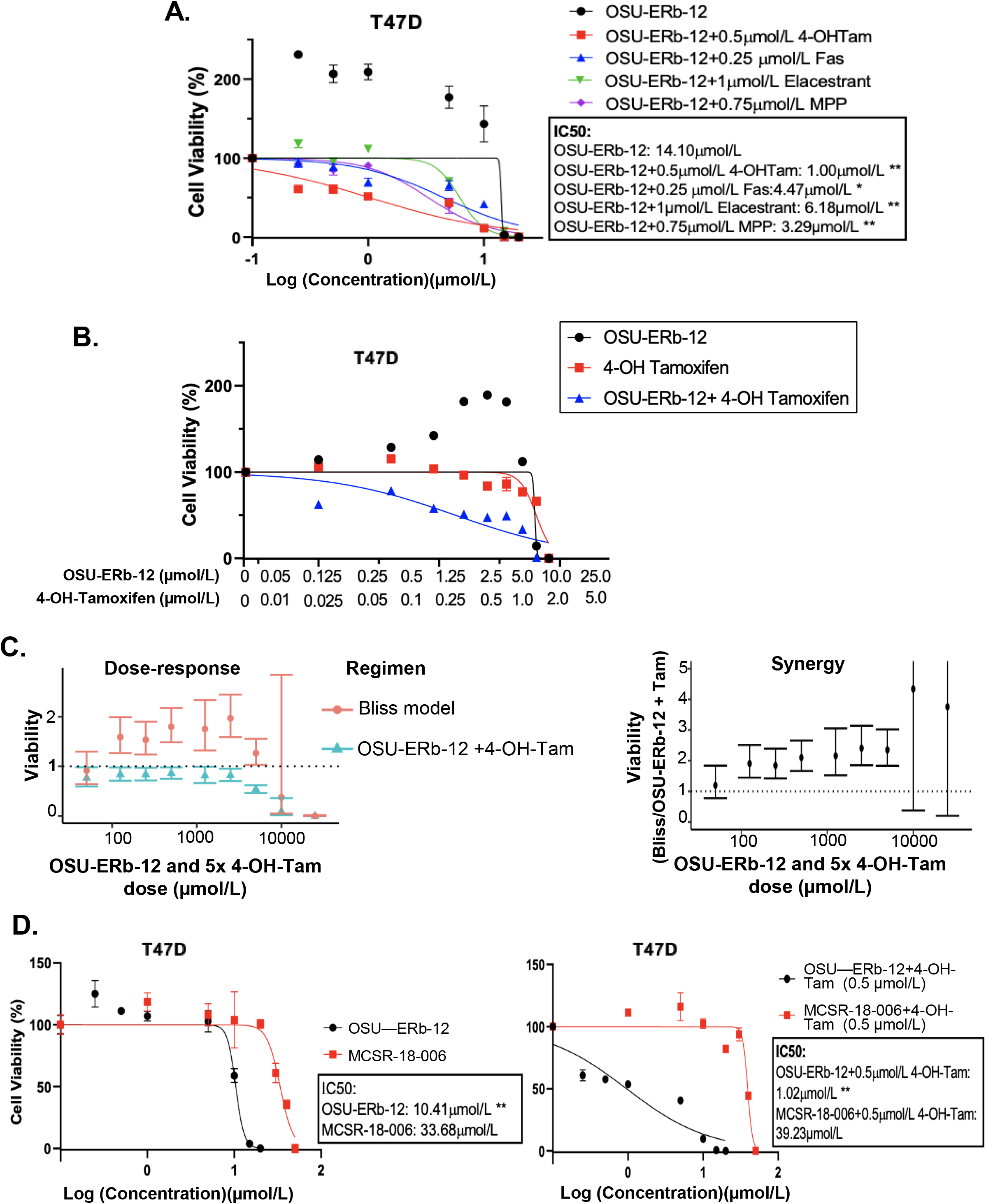
A-C, Combination treatment with selective ERβ agonists and ERα antagonist demonstrate significant cytotoxicity with reduction of IC50 in ERα+ breast cancer cell lines. **A**, T47D treated with: OSU-ERb-12 alone and combination with 4-hydroxy tamoxifen, fulvestrant, elacastrant, or MPP or **B**, OSU-ERb-12 alone, 4-hydroxy tamoxifen alone, and OSU-ERb-12 in combination with 4-hydroxy tamoxifen **C**, Linear mixed models were fit for viability versus regimen for each dose, with random effects accounting for within-batch correlation. Bliss independence model predictions are products of fitted values for 4-hydroxy tamoxifen and OSU-ERb-12. Error bars are 95% confidence intervals. (**left**); The ratio of predicted viabilities (Bliss independence / Combination 4-hydroxy tamoxifen + OSU-ERb-12) quantifies interaction, with ratios >1 indicating synergy. Error bars are 95% confidence intervals (**right**). **D**, T47D treated with: OSU-ERb-12 and MCSR-18-006 (**left**), combination of OSU-ERb-12/MCSR-18-006 with 4-hydroxy tamoxifen (**right**). Viable cells were measured after seven days of treatment with DMSO (control) or the drugs at the indicated concentrations using CellTiterGlo assay. The percentage of viable cells is shown relative to DMSO vehicle-treated controls (mean ± SD, *: p<0.05, **: p<0.01). Assays were performed in quadruplicates (three experimental replicates).

We next determined whether OSU-ERb-12 effects are specifically mediated by the ERβ receptor by comparing the OSU-ERb-12 induced decreases in cell viability to that of an inactive chemical analog MCSR-18-006 that differs at two atoms from OSU-ERb-12 (**Supplemental Fig. S2**). As shown in **Fig. 3D**, in T47D cells, OSU-ERb-12 demonstrated an IC50 value of 10.41 μmol/L that was 3.24-fold lower than for MCSR-18-006 (p<0.01). However, in the presence of 4-hydroxy-tamoxifen (0.5 μmol/L) the IC50 of OSU-ERb-12 was 1.02 μmol/L, which was 38.5-fold lower than that of MCSR-18-006 combined with 4-hydroxy-tamoxifen (**Fig. 3D, right figure; Supplemental Table 5 and 6)**.

We then tested the viability of both MCF7 and T47D cell lines upon treatment with three other less selective ERβ agonists namely, DPN (diarylpropionitrile) (15), AC186 (20), and WAY200070 (21). Our data demonstrated that none of these ERβ agonists (**Supplemental Fig. S6**) exerted any significant cytotoxic effect on any of the ERα+ cell lines.

### Selective ERβ agonists exert anti-proliferative and apoptotic effects on ERα+ breast cancer cell lines and results in induction of FOXO 1/3 proteins in ERα+ breast cancer cell lines

Since both the ERβ agonists reduced the viability of ERα+ cell lines we further examined the mechanism of reduced viability. Both OSU-ERb-12 and LY500307 reduced cell proliferation, induced S phase arrest and increased apoptosis of MCF7 and T47D cells (**Fig. 4**).

**Figure 4:**
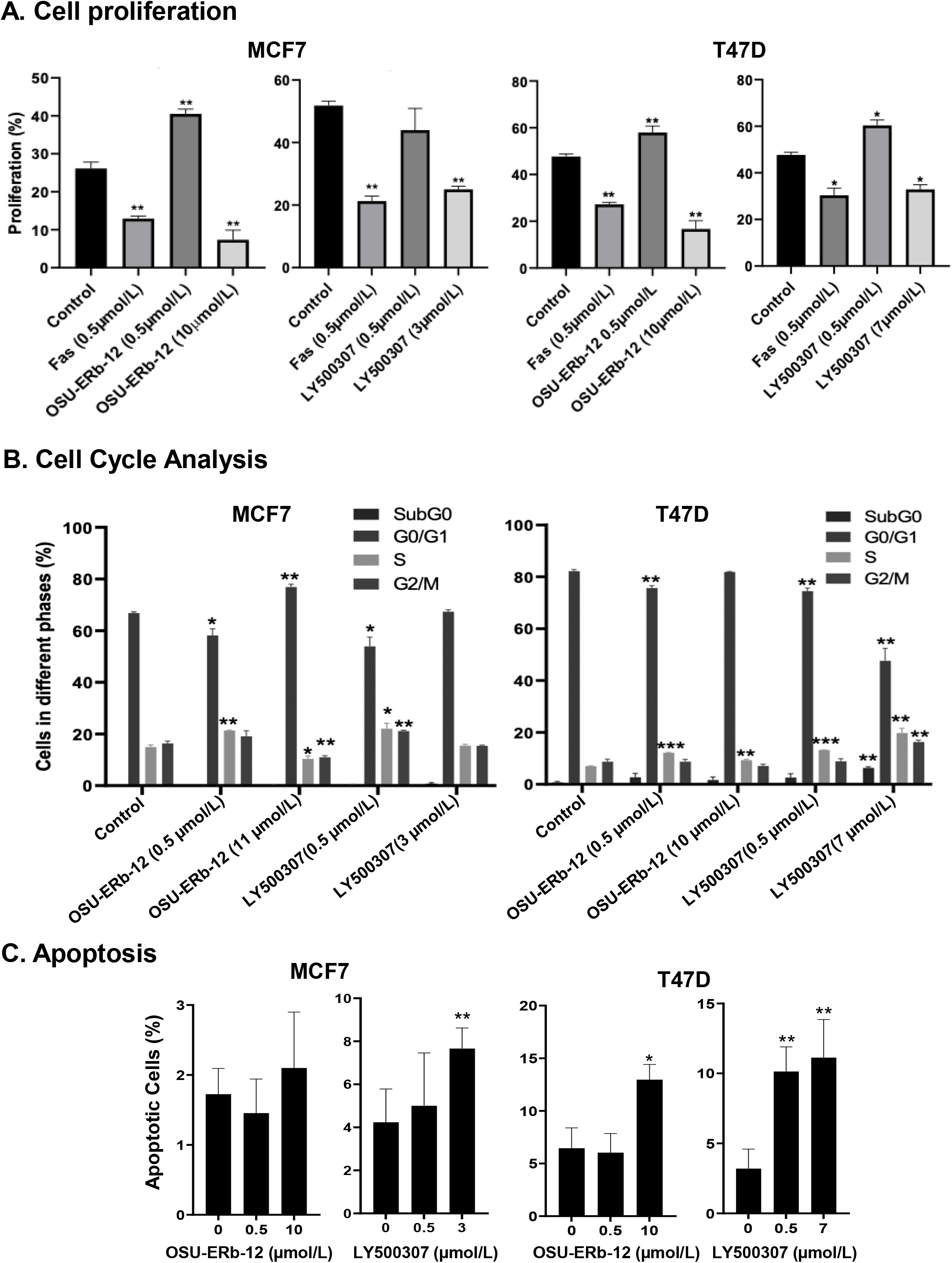
Cell proliferation, cell cycle and apoptosis are affected upon treatment of ERα+ breast cancer cells with ERβ specific agonists, OSU-ERb-12 and LY500307. MCF7 and T47D cells (0.5 × 10^6^) were seeded on 100 mm dishes in phenol red free DMEM containing charcoal stripped FBS and treated with the drugs as indicated. **A**, a representative diagram of cell proliferation profile in drug-treated cells. Cells were treated with DMSO (control), FAS (Fulvestrant; negative control), OSU-ERb-12 or LY500307 for 72 hours, harvested, and stained following protocol for the Click-iT Edu Alexa Fluor 647 kit (Invitrogen C10424). Cell proliferation was analyzed via flow cytometry on BD FACSCalibur Flow Cytometer. Each assay was performed in triplicate and repeated twice. Data were plotted as mean ± SD (*: p<0.05, **: p<0.01) **B**, a representative diagram depicting cell cycle profile in drug-treated cells. Cells treated with DMSO (control), OSU-ERb-12 or LY500307 for 72 hours at the indicated concentrations were harvested on ice, fixed, washed, and incubated with propidium iodide and RNase A followed by cell cycle analysis in a flow cytometer. Each assay was performed in triplicate and repeated twice. Data were plotted as mean ± SD (*: p<0.05, **: p<0.01, ***: p<0.001) **C**, a representative diagram depicting apoptosis profile in drug-treated cells. Cells treated with DMSO (control), OSU-ERb-12 or LY500307 for 48 hours at the indicated concentrations were harvested on ice, washed, and processed according to the manufacturer’s protocol (TUNEL Assay Kit-BrdU-Red; Abcam) followed by analysis on a BD FACSCalibur Flow Cytometer. Each experiment was repeated twice. Data presented are mean ± SD (*: p<0.05, **: p<0.01).

Cell proliferation was reduced by OSU-ERb-12 (10 μmol/L) and LY500307 (3 μmol/L) in MCF7 cells by 19% (p=0.016) and 27% (p=0.0028), respectively (**Fig.4A, Supplemental Fig. S7, Supplemental Table 7**). Similarly, in T47D cells OSU-ERb-12 (10 μmol/L) and LY500307 (7 μmol/L) reduced proliferation by 31% (p 0.0074) and 15% (p=0.015), respectively (**Fig.4A, Supplemental Fig. S7, Supplemental Table 7**). However, the observation that the ERβ agonists either significantly increased or did not decrease proliferation at the lower concentration (0.5 μmol/L) in both the cell lines, explains the increased cell viability observed at lower doses in earlier experiments (**Fig.2**).

Cell cycle analysis demonstrated that OSU-ERb-12 treatment (0.5 μmol/L) reduced the G0/G1 phase (8.7% decrease p=0.02) and increased S-phase fraction (6.4% increase, p=0.0347) of MCF7 as well as in T47D cells (G0/G1: 6.6% decrease, p=0.0036; S-phase: 5.2% increase, p= 0.0015) (**Fig. 4B, Supplemental Fig. S8, Supplemental Table 8)**. Similarly, LY500307 at 0.5 μmol/L caused a significant reduction in G0/G1 phase (13% decrease, p=0.019) and increase in S-phase (7.1% increase, p=0.049) of MCF7 as well as T47D cells (G0/G1: 7.7% decrease, p=0.0018; S-phase: 6.2% increase, p=0.0004) (**Fig. 4B, Supplemental Fig. S8, Supplemental Table 8)**. However, at a higher dose (around IC50) OSU-ERb-12 demonstrated no significant decrease in G0/G1 phase nor arrest at S -phase in both the cell lines-an observation that needs further explanation. Nevertheless, in T47D cells, LY500307 at higher dose (7 μmol/L) exhibited a dramatic decrease (34%, p=0.0079) of G0/G1 phase, increase in apoptotic cells (at SubG0, 5.6%, p=0.0068), arrest at S (12.8% increase, p=0.006), and G2/M (7.6% increase, p=0.0135) phases, respectively. Altogether, this data suggests that treatment with ERβ agonists causes cell cycle arrest in S and/or G2/M phases.

We observed a significant increase in apoptosis of LY500307-treated (7μmol/L) MCF-7 cells (7.7% apoptotic cells, p=0.01) compared to the vehicle-treated control (4.2% apoptotic cells). We did not observe a statistically significant increase in apoptosis in MCF7 cells treated with OSU-ERb-12. We noticed a significant increase in apoptosis of T47D cells treated with 10 μmol/L OSU-ERb-12 (13%, p=0.03), 0.5 μmol/L LY500307 (10.1%, p=0.003) and 7 μmol/L LY500307 (11.1%, p=0.0005) apoptotic cells, respectively as compared to the vehicle treated control (3.2%) (**Fig. 4C, Supplemental Fig. S9, Supplemental Table 9**).

Next, we tested the efficacy of OSU-ERb-12 and LY500307 in reducing colony formation of MCF7 and T47D cells. Colony-forming ability was significantly reduced upon treatment with both the agonists (**Fig. 5A, Supplemental Table 10)**. In comparison with vehicle-treated cells OSU-ERb-12 suppressed colony formation in MCF7 cells by 14% (p=0.05) and 44% (p=0.002) and LY500307 by 79% (p=0.003), and 100% (p=0.0007) at 3 μmol/L and 5 μmol/L, respectively. Similarly, the reduction in colony formation in T47D with OSU-ERb-12 was 64.5% (5 μmol/L; p=0.011). With LY500307 colony formation was reduced by 19.9% (3 μmol/L; p=0.015) and 95% (5 μmol/L; p=0.005). However, there was no significant reduction of colony formation in T47D treated with 3 μmol/L OSU-ERb-12 (**Fig. 5A, Supplemental Table 10)**.

**Figure 5:**
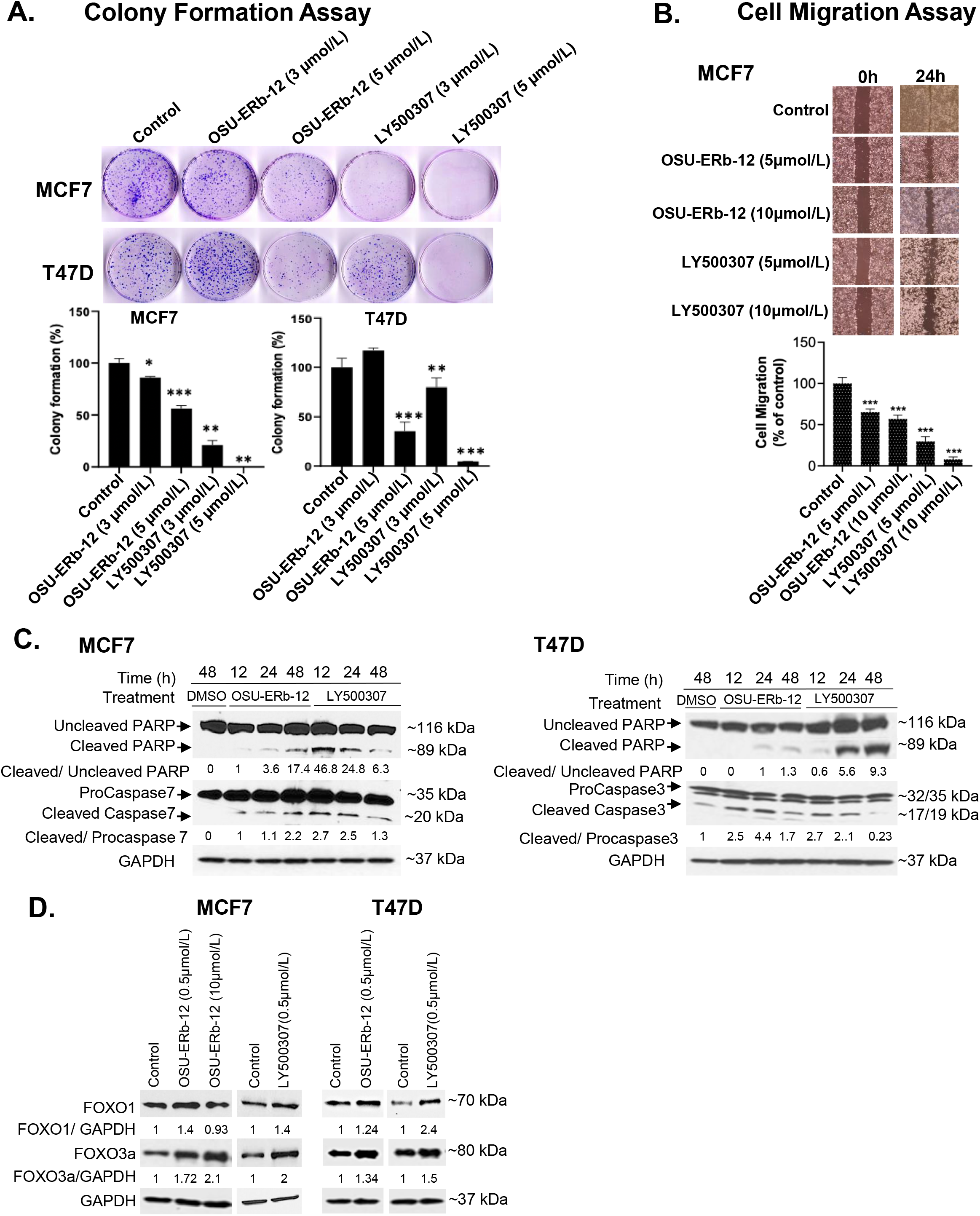
Treatment with ERβ specific agonists, OSU-ERb-129 and LY500307 promotes global anticancer effects in ERα+ breast cancer in vitro. **A**, colony formation. Colonies were stained with crystal violet and counted. The percentage of colonies present in each treatment is shown relative to DMSO vehicle-treated controls. Data are from three independent experiments and presented as mean ± SD; *: p < 0.05, **: p<0.01, ***: p<0.001; n = 3. **B**, cell migration. Cell migration was determined using the wound-healing assay. The percentage of filled area is calculated, normalized to DMSO treated control and presented as mean ± SD from three independent experiments; mean ± SD; *: p < 0.05, **: p<0.01, ***: p<0.001; n = 3. **C**, Enhanced cleavage of PARP-1, and activation of caspases 3 and 7 in ERα+ breast cancer cells upon treatment with ERβ agonists. Western blot analyses were performed using specific antibodies in whole cell lysates prepared from OSU-ERb-12 and LY500307 treated cells as indicated. Similar results were obtained in different batches of cells treated with OSU-ERb-12 and LY500307. Numbers under the lanes are quantitative representation of the intensity of the normalized bands. The signal in each band was quantified using Image Studio (LiCor) software. **D**, Enhanced expression of FOXO1 and FOXO3a proteins in ERα+ breast cancer cells upon treatment with ERβ agonists. Western blot analyses were performed using specific antibodies in whole cell lysates prepared from cells treated for 7 days with OSU-ERb-12 or LY500307. Similar results were obtained with different batches of cells treated with OSU-ERb-12 or LY500307. Numbers under the lanes represent corresponding normalized band intensity of the respective proteins. Image Studio (LiCor) software was used to quantify the intensity of the protein bands.

We then performed a cell motility assay to investigate whether OSU-ERb-12 and LY500307 treatment could lead to the reduction of migratory properties of breast cancer cells. As shown in **Fig. 5B**, there was a significant decrease in the cell motility in the MCF7 cell line in the presence of both the agonists. Treatment with OSU-ERb-12 inhibited MCF7 cell migration by 34.7 % (5 μmol/L; p=0.0004) and 42.9% (10 μmol/L; p=0.0026) and LY500307 by 70.2 % (5 μmol/L; p<0.0001) and 91.9% (10 <u>μmol/L</u>; p<0.0001) (**Fig. 5B, Supplemental Table 11**).

To elucidate the underlying mechanism of ERβ agonists-mediated cell death, we measured the levels of activated executioner caspases by Western blot analysis. As MCF7 cells do not express caspase 3 (22), we measured caspase 7 levels in this cell line. Robust activation of the effector caspases 7 (MCF7) or 3 (T47D) resulted within 12 hours of treatment of cells with both the agonists. The effect persisted at least up to 48 hours (**Fig. 5C)**. In contrast, in vehicle-treated cells increased caspase cleavage was not detected. A similar increase in the proteolysis of their substrate PARP-1 was noted in ERβ agonist-treated cells (**Fig. 5C)**.

It has been demonstrated that ERβ suppresses tumor growth and induces apoptosis by augmenting the transcription of the tumor suppressor genes *FOXO1* and *FOXO3* in prostate cancer (23). Therefore, we determined their expression levels in ERβ agonist-treated breast cancer cells. As shown in **Fig. 5D**, both FOXO1 and FOXO3a protein levels were increased in OSU-ERb-12- and LY500307-treated MCF7 and T47D cell lines.

### ERβ expression in human breast cancer samples

Previous studies suggested that distinct from ERα, ERβ inhibits transcription from promoters that incorporate estrogen response-tetradecanoyl phorbol ester (ERE-AP1) composite response elements (13). We hypothesized that the ERβ/*ESR2* mRNA expression levels in ERα+ human breast cancer samples would negatively correlate with those of genes with promoters that contain ERE-AP1 response elements and that there would be a positive association between *ESR2* mRNA expression levels and overall survival.

Thirty-seven patients with metastatic ERα+/HER2-breast cancer were included in this study. Demographic and clinical characteristics are displayed in **Supplemental Table 12**. All the patients in this cohort were female with a median age of 56 years (range 27-78). The patients were predominantly Caucasian (35, 95%) and most women were postmenopausal (23, 66%).

We found that the expression of the cyclin D1 gene, the classic target of estrogen-stimulated transcription through an AP1 response element, negatively correlated with that of ERβ/*ESR2* as measured using Spearman correlation coefficient (rho = -0.45, p = 0.005) (**Figure 6B**). ERβ/*ESR2* expression was also negatively correlated with that of ERα/*ESR1* (rho = -0.35, p = 0.033). However, ERβ/*ESR2* mRNA expression positively correlated with that of *IGFBP4* (rho = 0.58, p < 0.001) and *CXCL12* (rho = 0.54, p < 0.001) (**Fig. 6B**). The univariate Cox proportional hazards estimate for overall survival by *ESR2* expression was 0.54 (95% CI 0.06, 5.22), suggesting a positive trend that did not reach statistical significance in this numerically limited cohort (**Fig. 6A**).

**Figure 6.**
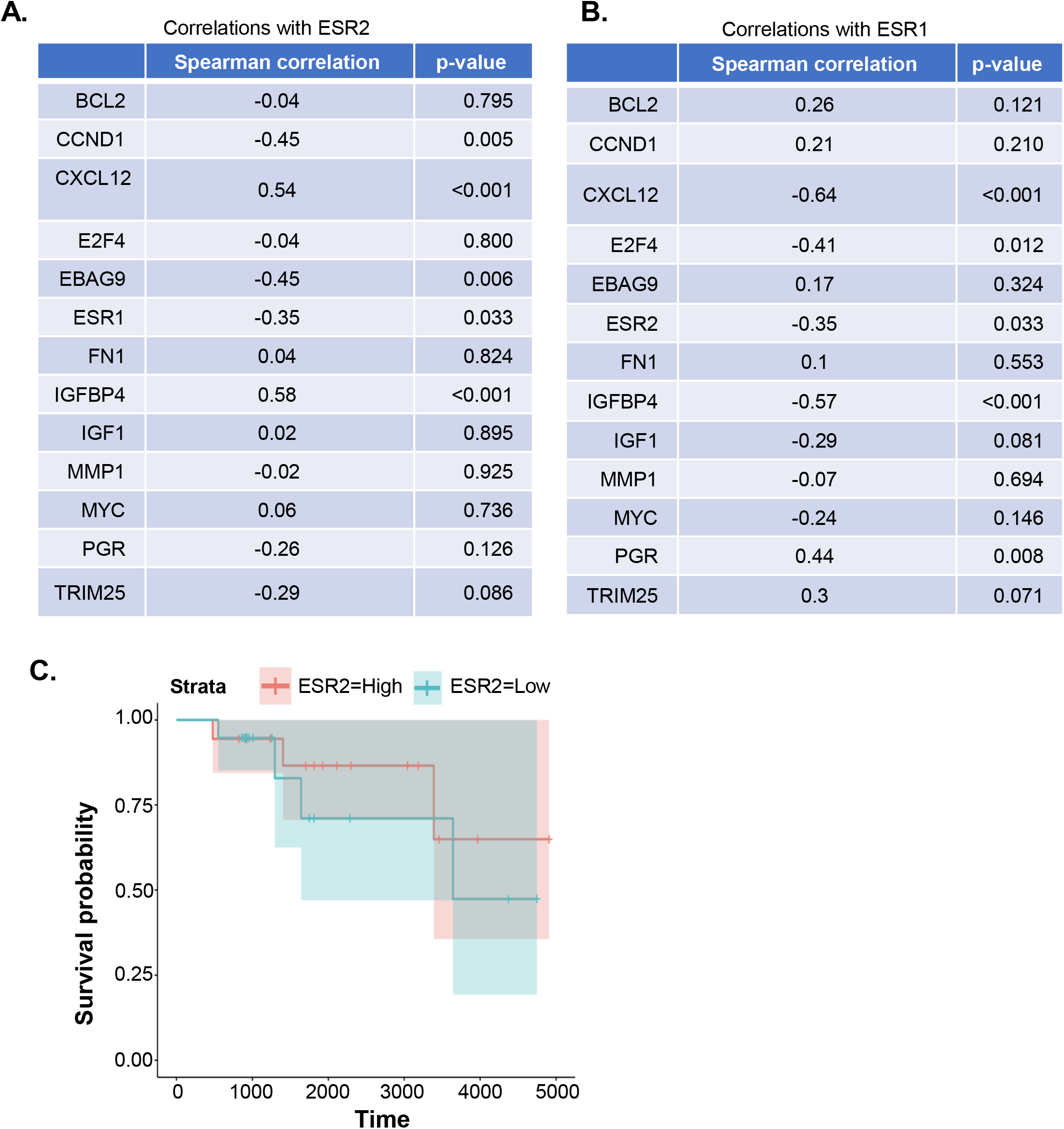
**A-B**, Expression of the genes that are targets of ER-AP1 mediated transcription and AP1 independent ERE mediated transcription in metastatic HER2 negative ER+ breast cancer patients is positively correlated with ESR2. A, ESR2 is positively correlated with CXCL12 and IGFBP4, and negatively correlated with CCND1, EBAG9, and ESR1, B, ESR1 is positively correlated with PGR and negatively correlated with CXCL12, E2F4, IGFBP4, and ESR2. Expression levels (FPKM), and Spearman correlation coefficients were computed for ESR1 and ESR2 versus other gene expression levels. **C**, Overall survival was not significantly correlated with the expression of ESR2 in the HER2 negative ERα+ metastatic breast cancer patient cohort, although there was a trend towards positive correlation. ESR2 was dichotomized relative to the median expression level and tested via the log-rank test (p = 0.6). Cox proportional hazards regression on the continuous expression levels yielded similar results (HR 0.6, p = 0.7).

## Discussion

ERα subtype constitutes 70% of all breast cancers while annually about 600,000 breast cancer-related death occurs worldwide (1). Although metastatic ERα+ breast cancer is initially treated with estrogen deprivation or ERα blockade, endocrine resistance eventually entails a change in therapy. The response to second-line endocrine agents such as fulvestrant is generally short. The advent of CDK4/6 inhibitors such as palbociclib (24, 25), ribociclib (26), and abemaciclib (27, 28) has doubled progression-free survival when used in combination with endocrine agents. However, resistance to CDK4/6 inhibitors is an increasing clinical challenge (29). Also, the duration of response to second-line endocrine therapies is generally short. After the exhaustion of endocrine treatment, chemotherapy remains the only treatment option. Therefore, there is an urgent need for tolerable therapies to prolong overall survival with better quality of life for advanced ERα+ breast cancer patients.

Accumulating evidence suggest while ERα is oncogenic, ERβ plays a tumor suppressor role in different cancers including breast cancer (30, 31). The efficacy of selective ERβ agonists such as LY500307 has been previously described in preclinical models of TNBC (32), melanoma (32), glioblastoma multiforme (33), and prostate cancer (34). However, there has been limited study of the role of ERβ in estrogen receptor α positive breast cancer. One reason is that for this particular indication a high degree of selectivity for ERβ over ERα would be required. Our institution recently developed a highly selective ERβ agonist: OSU-ERb-12 (16). We confirmed the selectivity of this compound using ERE-luciferase promoter assays showing ∼40-fold induction upon treatment of ERβ overexpressing cells.

Although previous preclinical studies have mostly focused on TNBC, we observed that ERβ was expressed (both RNA and protein level) in ERα+ breast cancer cell lines at levels that were not significantly different from those in TNBC cell lines (**Fig. 1, A-C)**. Endocrine and CDK4/6 resistant derivatives of these ERα+ cell lines had comparable or higher expression compared to the parental cell lines. These observations, therefore, are in line with the potential for efficacy in ERα+ breast cancer.

We showed that OSU-ERb-12, like the control compound LY500307, exerted significant cytotoxicity towards MCF7 and T47D ERα+ breast cancer cell lines with IC50 values were lower compared to immortalized mammary epithelial cells (MCF10A). Furthermore, OSU-ERb-12 exhibited cytotoxicity towards the corresponding endocrine- and CDK4/6 inhibitor-resistant derivative lines of MCF7 and T47D with either similar or even significantly lower IC50 values, demonstrating its therapeutic efficacy towards both treatment naïve and resistant ERα+ breast cancer cells. Furthermore, we demonstrated that these effects are ERβ specific using a close structural analog that lacks the ERβ agonist activity and was many-fold less cytotoxic than the active compound.

At lower concentrations of OSU-ERb-12 and LY500307, there was an increase in cell viability. We hypothesized that this may be due to ERα activation, given the large molar excess of ERα receptors over ERβ receptors in ERα+ breast cancer cell lines. This prompted us to investigate the cytotoxic efficacy of OSU-ERb-12 in combination with clinically available potent ERα antagonists. In the combination studies, tamoxifen showed maximum inhibitory effect with a 14-fold reduction of IC50 value compared with OSU-ERb-12 alone. Using the Bliss Independence model, we found synergistic interaction between tamoxifen and OSU-ERb-12 at all the doses used in the study.

Of note, the cellular 50% inhibitory concentration were many-fold higher than the cellular 50% effective concentration for activation of a canonical palindromic ERE response element. There are many potential explanations for this. Firstly, inhibition of viability may only be achieved when the majority of available receptor is activated by ligand, for example possibly at the EC90-100 concentration range. Secondly, the EC50 concentration represents transcriptional activation at a palindromic estrogen response element with optimal configuration and spacing of the half binding sites. Depending on the configuration of the EREs in promoters, the EC50 may be higher. Of note, ligand-ER-DNA interactions, including the stoichiometry and affinity of the ligand for the ligand-binding domain are dependent on the spacing and orientation of ERE binding sites as well as flanking sequences (35-37). Thirdly, cytotoxicity may not be dependent on transcription but on ligand-induced protein-protein interactions that may also modulate ligand binding (38).

Our study demonstrated the efficacy of ERβ agonists in attenuating cell proliferation, cell migration and colony formation as well as inducing cell cycle arrest and apoptosis of ERα+ breast cancer cell lines. Also, we showed that ERβ agonist treated MCF7 and T47D cells exhibited activation of effector caspases 7/3 and cleavage of PARP as well, which are markers of apoptosis. FOXO proteins act as tumor suppressors in a variety of cancers including breast cancer (39, 40). Previous studies have shown that ERβ upregulates the expression of FOXO transcription factors in preclinical models of prostate cancer (23, 41, 42). Our data demonstrated significantly higher expression of both FOXO1 and FOXO3a proteins in ERβ agonist-treated cells. Thus, induction of FOXO proteins may be one of the mechanism(s) by which OSU-ER-12 exhibits its tumor-suppressor activity. Further confirmation of the necessity of FOXO transcription factor upregulation for the efficacy of ERβ agonists will be required.

Given the tumor suppressor activity of ERβ, we hypothesized that its expression would be positively associated with the overall survival of metastatic breast cancer patients. In the present study, we showed that in a cohort of 37 metastatic breast cancer patients there was a trend of increased overall survival in *ESR2*-high expressing patients compared to *ESR2*-low expressing patients. However, this data is not statistically significant in this small cohort of patients. Further analysis in a larger cohort is warranted. Previous studies had suggested that ERβ might antagonize the transcriptional upregulation of genes that incorporate composite estrogen-phorbol ester response elements such as *CCND1* (43-45). In our cohort of patients, we found that the expression of *CCND1* mRNA, a typical estrogen-stimulated target gene, is negatively correlated with the expression of *ESR2* mRNA.

In conclusion, we have provided sufficient evidence that OSU-ERb-12 could be a potential candidate compound for its tumor suppressor activity towards ERα+ breast cancer. Understanding the details of its mechanism of action and further confirmation of its efficacy is warranted using *in vivo* model systems.

## Author Contributions

MAC and JD conceived the project. JD, CCC, BR, and MAC designed the experiments. BR, ML, DGS, SDS, and MAC helped recruit the patients to the protocol under which the patient data were collected. JD, NW, JMM, PS, MS, JJD, DGS, and MK performed the experiments and analyzed the data. JD and MAC wrote the manuscript and all other authors reviewed and edited the manuscript.

## Acknowledgements

We would like to like to thank the Comprehensive Cancer Center, Arthur G. James Cancer Hospital and Richard Solove Research Institute at the Ohio State University Wexner Medical Center for supporting the study. We also would like to acknowledge Jackie Sharpnack for administrative support.

