## Supplemental Data for "Activity of Estrogen Receptor β Agonists in Therapy-Resistant Estrogen Receptor-Positive Breast Cancer"

#### Preparation of MCSR-18-006

#### Synthetic Scheme for the Preparation of MCSR-18-006

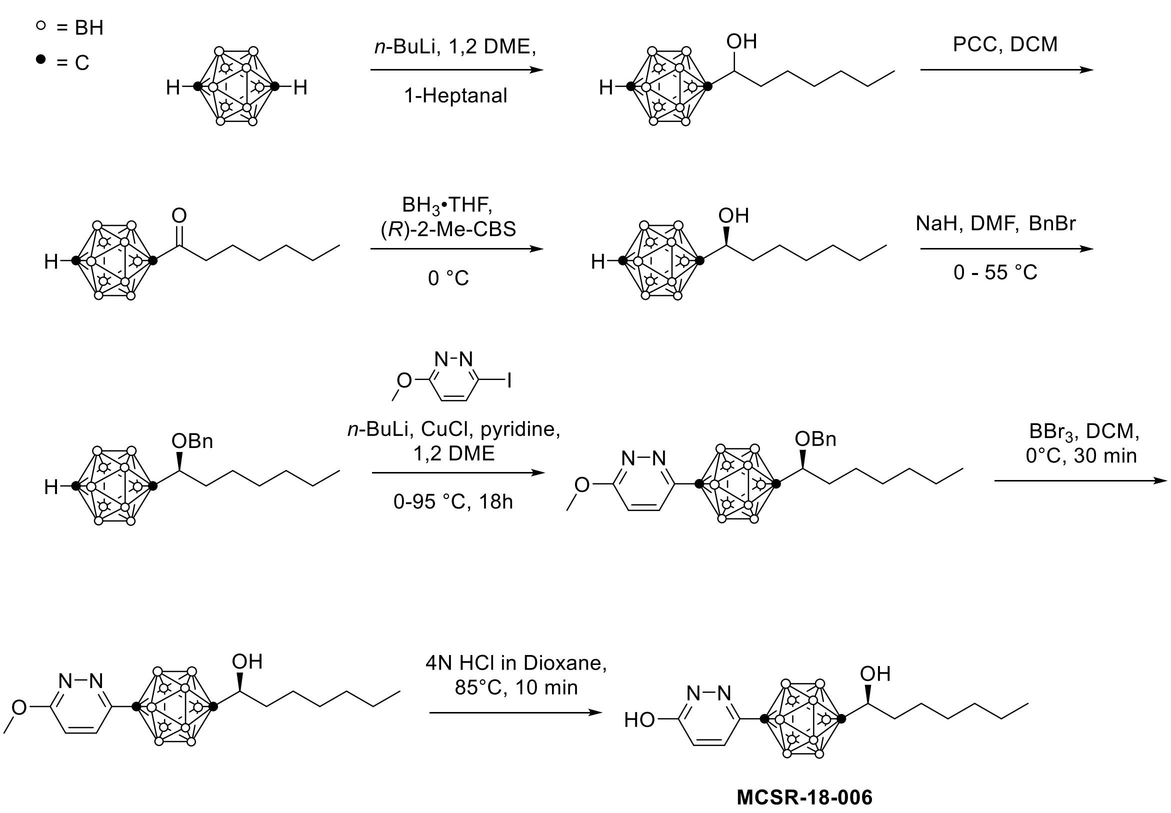

#### Procedure for the synthesis of MCSR-18-006

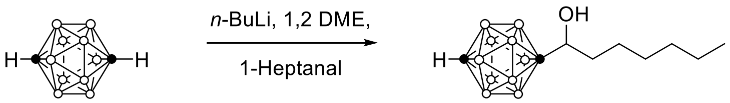

*1-(1-hydroxyheptyl)- 1,12-dicarba-closo-dodecaborane:* *n*-BuLi solution (2.5M in hexane, 4.4 ml) was added dropwise at 0 °C under Ar to a solution of 1,12-dicarba-*closo*-dodecaborane (1.44 g, 10 mmol) in 1,2 dimethoxyethane (50 ml) . We stirred the mixture at room temperature for 1 hour and then added 1-heptanal (1.55 ml, 11 mmol) at 0 ºC. This mixture was then stirred at room temperature overnight.  We then poured the mixture into 1 M HCl aqueous solution (100 ml), and performed extraction with ethyl acetate (3 X 25 ml). We then washed the combined organic phases with brine and dried them over $MgSO_4$. We evaporated the solvents and purified the residue using Teledyne Isco (RediSepRf column) to yield a colorless oil. Yield 1.4g. $^1$H NMR ($CDCl_3$) δ 3.38-3.45(m, 1 H), 3.35-1.14 (m, 22H), 0.88 (t, 3 H), MS 258.291.

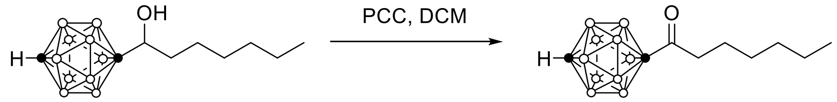

*1-(Heptan-1-one)-1,12-dicarba-closo-dodecaborane:* We suspended pyridinium chlorochromate (PCC, 1.7 g, 7.71 mmol) in anhydrous DCM (50 ml). We then added a solution of 1-(heptan-1-yl)-1,12-dicarba-*closo*-dodecaborane (1.3 g, 5.04 mmol) in DCM (10 ml), and stirred the reaction mixture at room temperature overnight. Diethyl ether (50 ml) was added to the mixture followed by molecular sieves, and then stirred for 1 h. We decanted the supernatant and washed the insoluble residue with dry ether (3 x 20 ml). We then passed the combined organic phases through a short column of Celite followed by evaporation. We then purified the residue with Teledyne Isco (RediSepRf column) to yield a colorless oil, yield 1.2g. $^1$H NMR ($CDCl_3$) δ 1.16 (m, 21H), 0.88 (t, 3 H), MS 256.189.

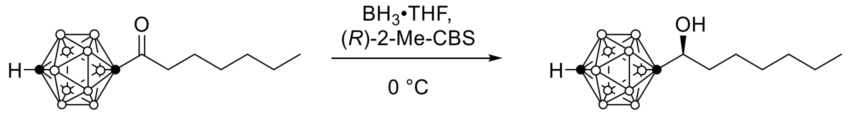

*(R)-1-(1-hydroxyheptyl)- 1,12-dicarba-closo-dodecaborane:* We added Borane-tetrahydrofuran complex (51 mL, 51 mmol, 1.0 M solution in THF, stabilized with 0.005 M N-isopropyl-N-methyl-tert-butylamine (NIMBA) followed by (R)-2-methyl-CBS- oxazaborolidine [(2-MeCBS] (5.1 mL, 5.1 mmol, 1.0 M solution in toluene) to 50 mL anhydrous THF. The reaction mixture was stirred at room temperature for 15 minutes. We then added 1-(Heptan-1-one)-1,12-dicarba-

*closo*-dodecaborane (1.3 g, 5.08 mmol) in 25 mL of anhydrous THF slowly over a period of 2 h at 0 °C. The reaction mixture was stirred overnight at room temperature. We then carefully quenched the mixture by the addition of 2.0 M HCl (80 mL) in small portions to control $H_2$ development. We added diethyl ether (100 mL) and washed the organic phase with brine and saturated $NaHC0_3$. We then dried the organic phase over $MgS0_4$, filtered, and evaporated. The residue was then purified by Teledyne Isco (RediSepRf column) to yield a colorless oil, yield 1.1g 81%. $^1$H NMR ($CDCl_3$) δ 3.38-3.45(m, 1 H), 3.35-1.14 (m, 22H), 0.88 (t, 3 H), MS 258.291.

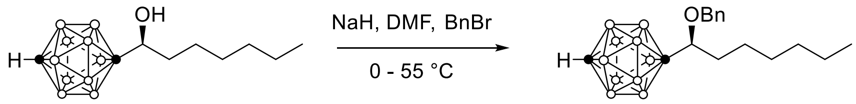

*(R)-1-(1-benzyloxy)heptyl)- 1,12-dicarba-closo-dodecaborane*: We added NaH (60% in mineral, 175 mg, 4.36 mmol) in one portion at 0 ºC to a solution of *(R)*-1-(1-hydroxyheptyl)- 1,12-dicarba-*closo*-dodecaborane (900 mg, 3.49 mmol) in anhydrous DMF (10 ml), , and then stirred at same temperature for 30 min. We then added BnBr (746 mg, 4.36 mmol) and stirred the reaction mixture at 55°C for 3 h. We then cooled it down to room temperature and added methanol (0.5 ml) slowly, followed by dilution with ethyl acetate (50ml), washing with water and brine and drying with $Na_2SO_4$. The solvents were then evaporated and the residue purified by Teledyne Isco (RediSepRf column) to yield a colorless oil, yield 1.1g 93%. $^1$H NMR (300 MHz, $CDCl_3$) δ 7.28-7.35 (m, 5H), 4.63 (d, *J* = 11 Hz), 4.30 (d, *J* = 11 Hz), 3.12-3.23 (m, 1H), 2.75 (s, 1H), 0.98-1.40 (m, 10H), 0.96-3.30 (m, 10H), 0.87 (t, *J* = 7 Hz, 3H).

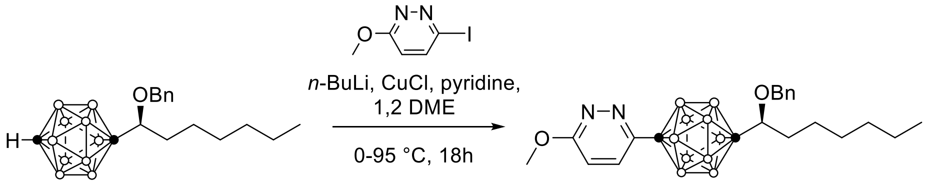

*(R)-1-(1-(benzyloxy)heptyl)-12-(6-methoxypyridazin-3-yl)-1,12-dicarba-closo-dodecaborane: n-*BuLi solution (2.5 M in hexanes, 274 µl, 0.68 mmol) at 0°C under Ar was added dropwise to a solution of (*R*)-1-(1-benzyloxy)heptyl)- 1,12-dicarba-*closo*-dodecaborane (197 mg, 0.56 mmol) in 1,2-dimethoxyethane (14.1 ml) . We then stirred the mixture at room temperature for 1h, and added CuCl (68 mg, 0.68 mmol) in one portion. Stirring was continued at room temperature for 1 h. We then added pyridine (637 µl, 7.91 mmol), and 3-iodo-6-methoxypyridazine (161 mg, 0.68 mmol) sequentially and heated the mixture at 95°C for 18 h. Following cooling, the reaction mixture was diluted with $Et_2O$ and then stirred at room temperature for 3 h. We then filtered the insoluble materials through a short pad of silica, washed the filtrate with 10% aqueous $Na_2S_2O_3$ and brine, and dried the organic layer over $Na_2SO_4$, followed by filtration and concentration. We then purified the resulting residue using Combiflash ($SiO_2$, RediSepRf 12 g gold column, gradient: EtOAc/Hex) to yield 108 mg (42%) of a clear colorless oil. $^1$H NMR (300 MHz, $CDCl_3$): δ 7.22-7.38 (m, 6H), 6.80 (d, *J* = 9.2 Hz, 1H), 4.65 (d, *J* = 11.0 Hz, 1H), 4.33 (d, *J* = 11.0 Hz, 1H), 4.09 (s, 3H), 3.23-3.29 (m, 1H), 1.41-3.66 (m, 10H), 1.01-1.44 (m, 10H), 0.85 (t, *J* = 7.0 Hz, 3H).

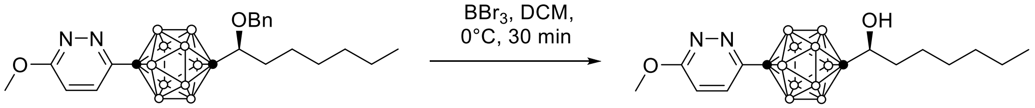

*(R)-1-(1-hydroxyheptyl)-12-(6-methoxypyridazin-3-yl)-1,12-dicarba-closo-dodecaborane:* 1 M solution of $BBr_3$ in $CH_2Cl_2$ (0.70 mL, 0.70 mmol) was added dropwise at 0 °C to a solution of *(R)*-1-(1-(benzyloxy)heptyl)-12-(6-methoxypyridazin-3-yl)-1,12-dicarba-*closo*-dodecaborane (91 mg, 0.20 mmol) in $CH_2Cl_2$ (2 ml). The mixture was stirred at 0 °C for 30 min and then poured into ice water and $NaHCO_3$. We then extracted the mixture 3x with $CH_2Cl_2$ (*note: 1-2% methanol was added to clarify the emulsion*). We washed the organic layer with brine, dried it over $Na_2SO_4$, and filtered and concentrated it *in vacuo*. We purified the resulting residue using Combiflash ($SiO_2$, RediSepRf 4g column, gradient: EtOAc/Hex) to yield 61 mg (83%) of a white powder. $^1$H NMR (400 MHz, $CDCl_3$): δ 7.25 (d, *J* = 9.3 Hz, 1H), 6.80 (d, *J* = 9.3 Hz, 1H), 4.09 (s, 3H), 3.44-3.50 (m, 1H), 1.63-3.40 (m, 10H), 1.59 (d, *J* = 6.2 Hz, 1H), 1.11-1.49 (m, 10H), 0.87 (t, *J* = 6.9 Hz, 3H).

**MCSR-18-006**

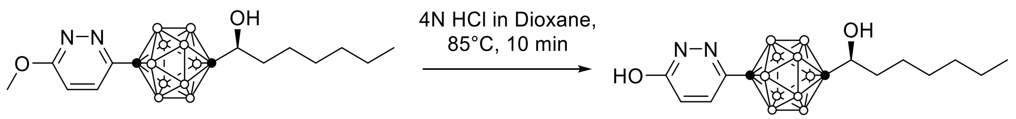

*(R)-1-(1-hydroxyheptyl)-12-(6-hydroxypyridazin-3-yl)-1,12-dicarba-closo-dodecaborane* : We added a solution of 4N HCl in Dioxane (0.5 mL) to *(R)*-1-(1-hydroxyheptyl)-12-(6-methoxypyridazin-3-yl)-1,12-dicarba-*closo*-dodecaborane (50 mg, 0.14 mmol). We then heated the mixture at 85 °C for 10 min, cooled it to room temperature, evaporated it to dryness with a gentle stream of argon, treated the residue with saturated aq. $NaHCO_3$ and extracted it 3x with ethyl acetate. We then washed the organic layer with brine, dried it over $Na_2SO_4$, filtered and concentrated it *in vacuo*. We purified the resulting residue using Combiflash ($SiO_2$, RediSepRf 4g gold column, gradient: MeOH/DCM) to yield 38 mg (79%) of a white powder. $^1$H NMR (400 MHz, $CDCl_3$): δ 10.79 (br s, 1H), 7.14 (d, *J* = 10.0 Hz, 1H), 6.79 (d, *J* = 10.0 Hz, 1H), 3.40-3.50 (m, 1H), 1.66-3.30 (m, 10H), 1.62 (d, *J* = 6.3 Hz, 1H), 1.11-1.50 (m, 10H), 0.87 (t, *J* = 6.9 Hz, 3H). MS (APCI-): *m/z* calcd. for $C_{13}H_{27}B_{10}N_2O_2$ $(M-H)^-$ 351.3, found 351.1.

#### NMR Spectra for carborane compounds

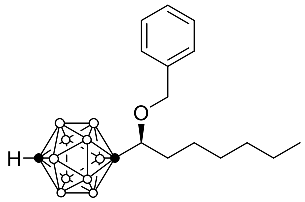

Molecular Weight: 348.53
Chemical Formula: $C_{16}H_{32}B_{10}O$

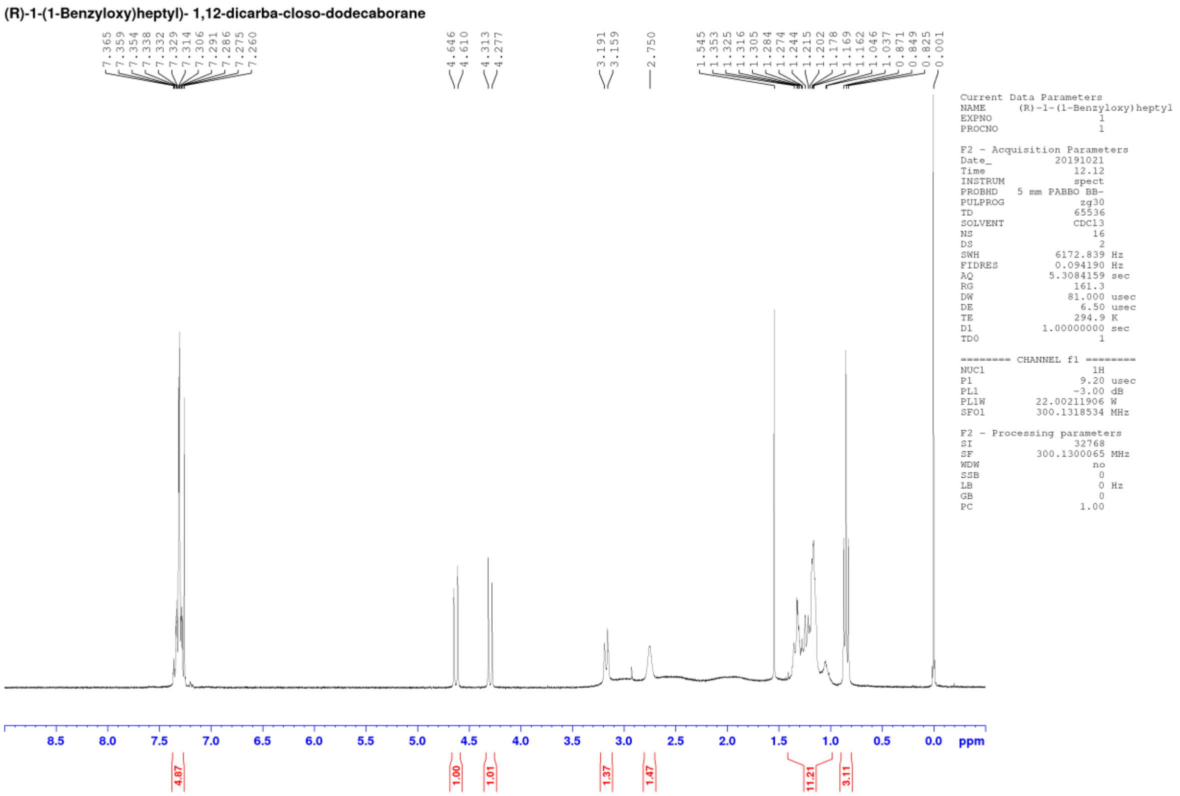

[1]H NMR ($CDCl_3$, 300MHz) of Compound *(R)*-1-(1-benzyloxy)heptyl)- 1,12-dicarba-*closo*-dodecaborane.

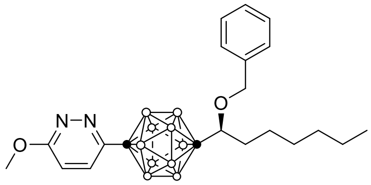

Molecular Weight: 456.63
Chemical Formula: $C_{21}H_{36}B_{10}N_2O_2$

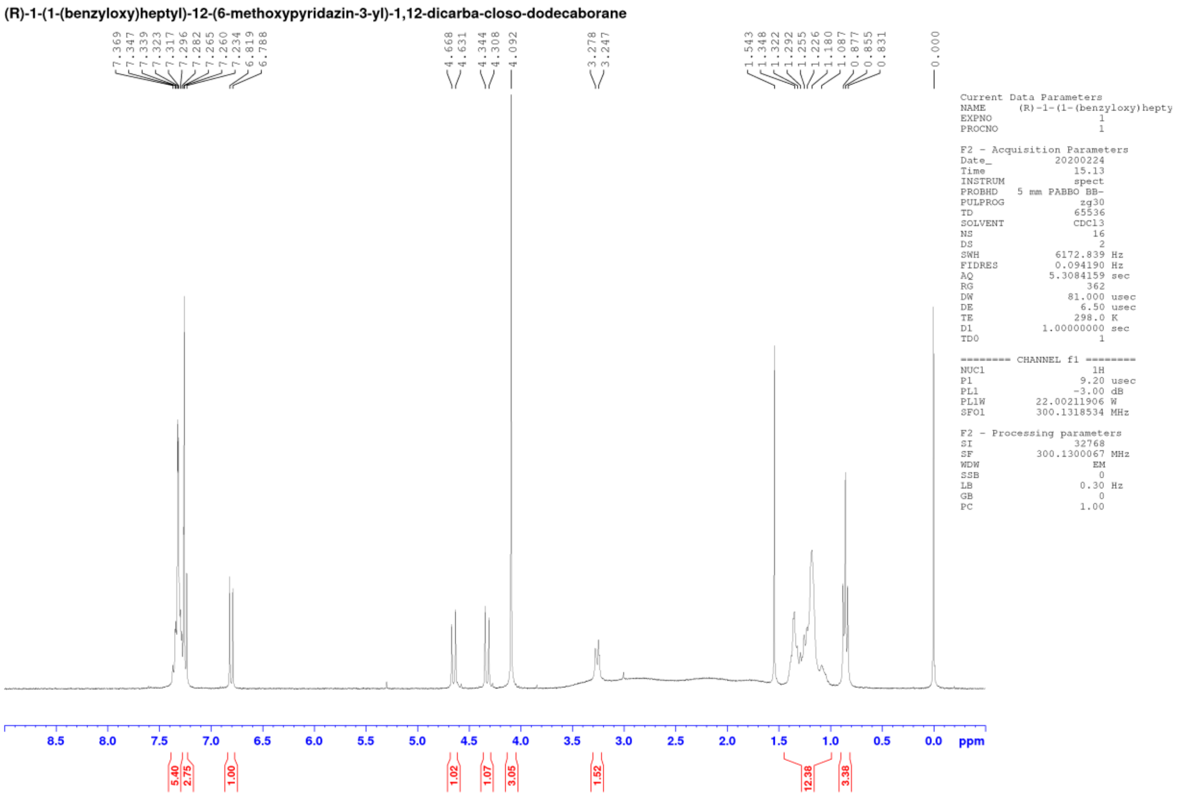

[1]H NMR (CDCl$_3$, 300MHz) of Compound *(R)*-1-(1-(benzyloxy)heptyl)-12-(6-methoxypyridazin-3-yl)-1,12-dicarba-*closo*-dodecaborane.

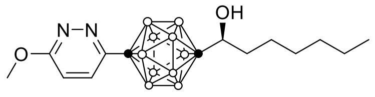

Molecular Weight: 366.51

Chemical Formula: $C_{14}H_{30}B_{10}N_2O_2$

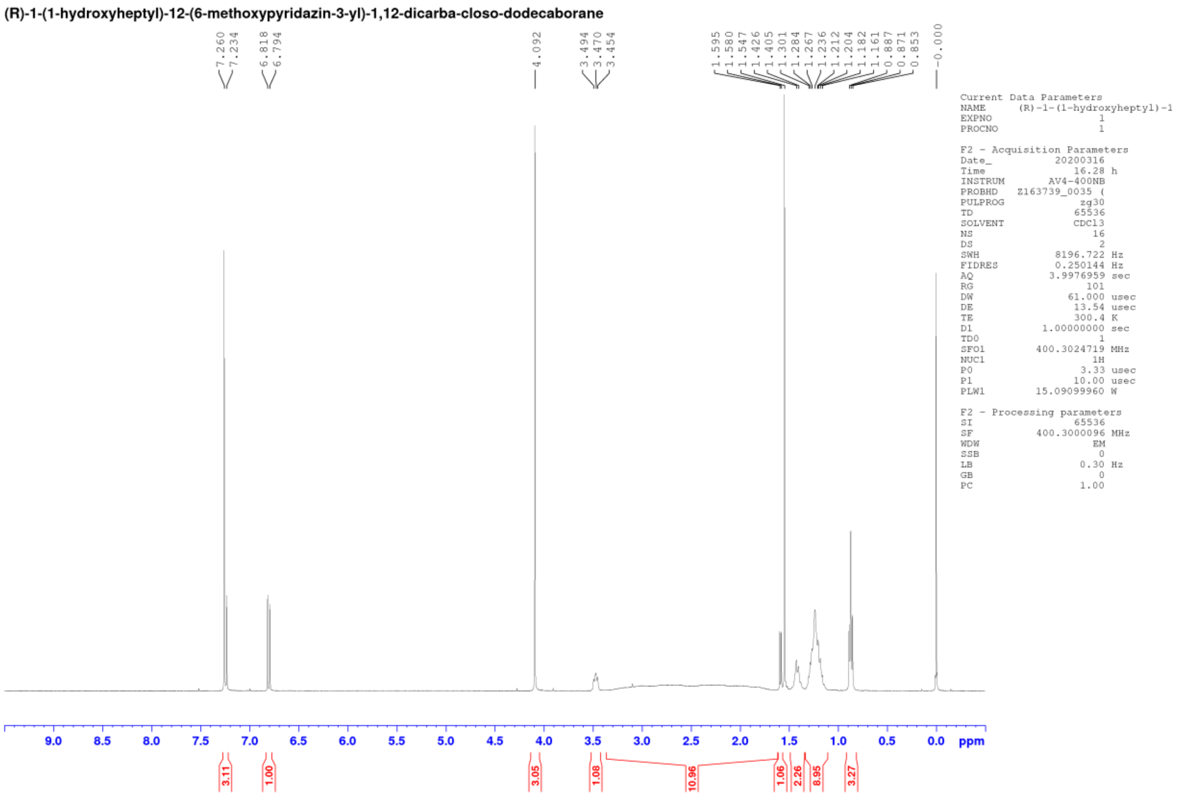

[1]H NMR ($CDCl_3$, 400MHz) of Compound *(R)*-1-(1-hydroxyheptyl)-12-(6-methoxypyridazin-3-yl)-1,12-dicarba-*closo*-dodecaborane.

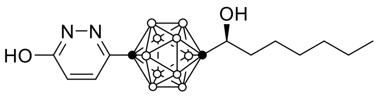

Molecular Weight: 352.48
Chemical Formula: $C_{13}H_{28}B_{10}N_2O_2$

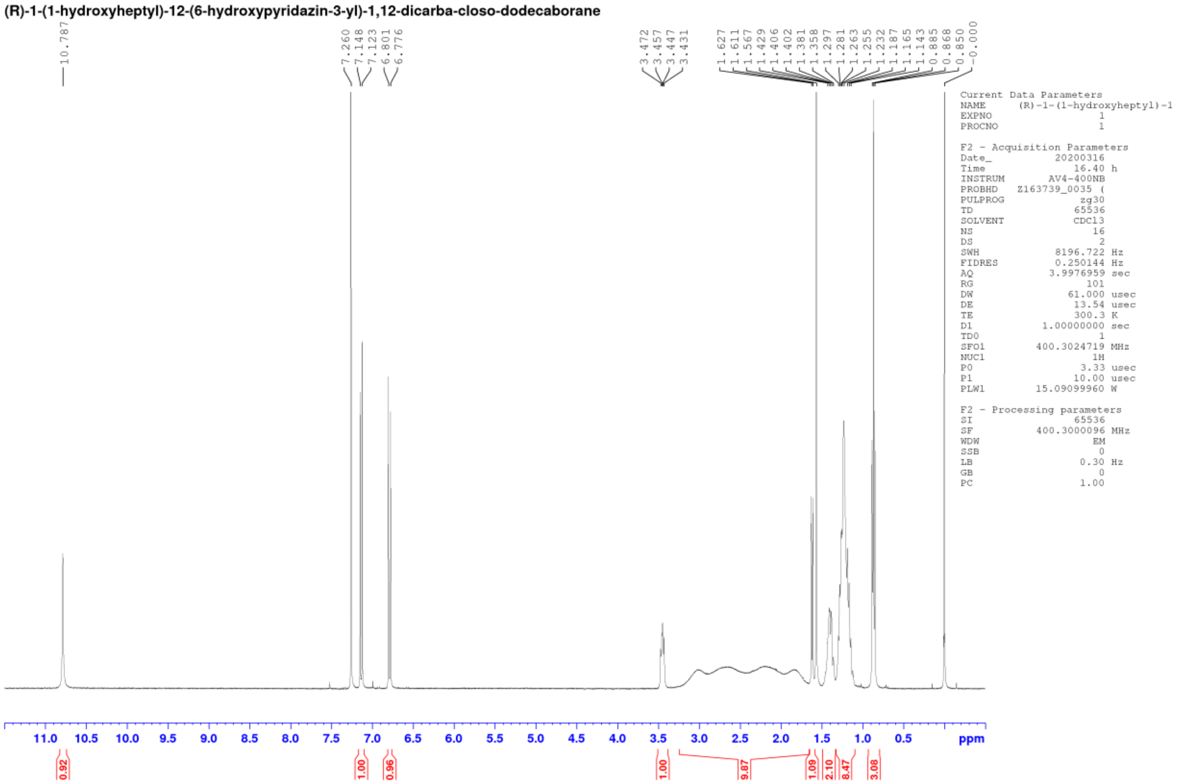

[1]H NMR ($CDCl_3$, 400MHz) of Compound *(R)*-1-(1-hydroxyheptyl)-12-(6-hydroxypyridazin-3-yl)-1,12-dicarba-*closo*-dodecaborane (**MCSR-18-006**).

**Supplemental Figure legends**

**Supplemental Figure S1.** Development of resistance in ERα+ breast cancer cells after chronic exposure to estrogen receptor antagonists and CDK4/6 inhibitor. Resistance was induced by chronic exposure of MCF7 and T47D cell lines to estrogen receptor antagonists and CDK4/6 inhibitor**.** Control cells were treated with DMSO in parallel. Control and resistant cells were treated for 72 hours with different concentrations of the drugs as indicated and the cell viability was measured. The percentage of viable cells relative to DMSO (control) are shown as mean ± SD. All the assays were performed in quadruplicate and repeated twice. Treatment with: **A,** 4-OH-tamoxifen **B,** fulvestrant **C,** abemaciclib.

**Supplemental Figure S2.** Chemical Structures of the inhibitors used in the study

**Supplemental Figure S3.** ERα/ERβ driven ERE-Luciferase promoter activity is unaffected by MCSR-18-006, the chemical analog of the ERβ specific agonist OSU-ERb-12. The cell viability was measured in extracts prepared from human full-length ERα or ERβ expressing Sf9 cells in presence of vehicle or different concentrations of estrogen (E2) or MCSR-18-006 as indicated. Fold change in cell viability was determined by comparing with the vehicle control.

**Supplemental Figure S4.** Radio-ligand displacement assay demonstrated insignificant binding of the non-specific chemical analog MCSR-18-006, to ERα/ERβ. Recombinant human full-length ERα and ERβ were expressed in Sf9 cells, prepared the cell lysates, and the relative binding affinity of MCSR-18-006 was measured by radiolabeled $^3$H estradiol competition binding assays. The IC50 the reference compound diethylstilbestrol was more than 100-fold less compared to MCSR-18-006 towards both ERα and ERβ. The binding affinities (Kis) of the reference compound for ERα and ERβ were 0.38 nmol/L and 0.22 nmol/L. The binding affinity value of MCSR-18-006 for ERα or ERβ could not be determined.

**Supplemental Figure S5.** **A,** Schematic diagram of primer design for qRT-PCR amplification of full length and all isoforms of *ESR2* gene. **B,** Agarose gel electrophoresis of qRT-PCR products of ESR2 full length, all isoforms, ESR1, and GAPDH showing a single band (of projected length) of corresponding gene product demonstrated specificity of the primers used in the study. qRT-

PCR products of the indicated genes with the RNA isolated from the cell lines were separated on 2% agarose gels, stained with Gel Green dye, and photographed. Each assay was done in triplicate and repeated twice. Similar results were obtained. A representative picture of each gene product has been shown.

**Supplemental Figure S6.** Less selective ERβ agonists did not reveal cytotoxicity in ERα+ breast cancer cell lines. Cell viability assays were performed after 7 days of treatment of MCF7 and T47D cells with the agonists. Cell viability was determined relative to the viable cells upon treatment with the vehicle control (DMSO). Each assay was performed in quadruplicate and repeated twice. Data was presented as mean ± SD. Similar results were obtained. Representative data has been presented. Treatment with: **A,** DPN **B,** AC186 **C,** WAY200070.

**Supplemental Figure S7.** Specific ERβ agonists OSU-ERb-12 and LY500307 inhibited proliferation of ERα+ breast cancer cells. Cell proliferation was measured after treatment with DMSO (control), fulvestrant, OSU-ERb-12 or LY500307 for 3 days. Washed cells were processed and proliferation was measured by FACS analysis as described in detail before (Materials and Methods). Each experiment was done in triplicate and repeated twice. Similar results were obtained. A representative profile has been presented. **A,** MCF7 **B,** T47D.

**Supplemental Figure S8.** Specific ERβ agonists OSU-ERb-12 and LY500307 caused S and/or G2/M phase arrest in ERα+ breast cancer cells. Cell cycle analyses were performed after treatment of cells with DMSO (control), OSU-ERb-12 or LY500307 for 3 days. Washed cells were processed and cell cycle profile was measured by FACS analysis as described in detail before (Materials and Methods). Each experiment was done in triplicate and repeated twice. Similar results were obtained. A representative profile has been presented. **A,** MCF7 **B,** T47D.

**Supplemental Figure S9.** Specific ERβ agonists OSU-ERb-12 and LY500307 induced apoptosis in ERα+ breast cancer cells. Apoptosis was measured after treatment of cells with DMSO (control), OSU-ERb-12 or LY500307 for 48 hours. Washed cells were processed, stained and apoptosis was measured by FACS analysis as described in detail before (Materials and Methods).

Each experiment was done in triplicate and repeated twice. Similar results were obtained. A representative profile has been presented. **Top,** MCF7 **Bottom,** T47D.

**Supplemental Table 1.** P-values and 95% confidence intervals (CIs) were determined for expression of the indicated genes in each cell line compared with that in the normal mammary epithelial line MCF10A.

| qRT-PCR | | | | | | |
|---|---|---|---|---|---|---|
| | ESR2 Full Length | | ESR2 All Isoforms | | ESR1 | |
| **Cell Line** | **p-value** | **95% CI** | **p-value** | **95% CI** | **p-value** | **95% CI** |
| MCF10A | | 0.862-1.420 | | 0.994-1.131 | | 0.664-1.408 |
| MCF7 | 0.7665 | 0.962-1.440 | 0.0524 | 2.999-3.998 | 0.0311 | 49770798-6034.474 |
| MCF7-TamR | 0.0035 | 3.240-4.291 | 0.0106 | 24.990-37.228 | 0.0179 | 9606.865-10728.515 |
| MCF7-FasR | 0.1196 | 1.502-1.526 | 0.0369 | 1.533-2.193 | 0.0227 | 429.422-493.742 |
| MCF7-CDK6 O/E | 0.1411 | 1.797-3.375 | 0.001 | 17.438-23.868 | 0.0147 | 2616.120-2864.450 |
| MCF7-CDK4/6iR | 0.0001 | 5.051-7.505 | 0.0018 | 31.568-54.566 | 0.0041 | 286.422-294.303 |
| T47D | 0.0265 | 6.536-8.202 | 0.001 | 72.211-81.421 | 0.0425 | 11597.865-15091.175 |
| T47D-TamR | 0.0009 | 4.698-5.294 | 0.0197 | 10.835-19.461 | 0.0203 | 3880.165-4398.077 |
| T47D-FasR | 0.0617 | 3.377-5.023 | 0.0194 | 26.819-46.711 | 0.0072 | 3071.065-3209.830 |
| T47D-CDK4/6iR | 0.0075 | 4.531-5.427 | 0.002 | 25.766-30.575 | 0.0331 | 7660.326-9398.086 |
| ZR-75-1 | 0.0001 | 19.319-20.873 | 0.0145 | 201.031-219.808 | 0.052 | 230.515-318.321 |
| MDA-MB-231 | 0.0404 | 3.616-4.985 | 0.0075 | 10.580-14.571 | 0.0496 | 1.577-1.896 |
| MDA-MB-468 | 0.0003 | 5.041-5.283 | 0.0761 | 3.191-4.695 | 0.0158 | 76.036-84.139 |

**Supplemental Table 2.** 95% confidence intervals (CIs) and p-values were determined for ERE-Luciferase activity in ERα and ERβ overexpressing cells treated with the ERβ agonists OSU-ERb-12 and LY500307. Left & middle columns: p-values were determined for each protein compared with corresponding untreated control. Right column: p-values were determined for each concentration of agonist treated ERβ overexpressing cells compared with that of ERα overexpressing cells.

| ERE-Luciferase Activity | | | | | |
|---|---|---|---|---|---|
| **OSU-ERb-12** | | | | | |
| | **ERα** | | **ERβ** | | **ERα vs ERβ** |
| **Concentration (µmol/L)** | **p-value** | **95% CI** | **p-value** | **95% CI** | **p-value** |
| 0 | | 0.92-1.08 | | 0.83-1.17 | 1 |
| 0.01 | 0.1435 | 0.8296-0.9896 | 0.034 | 2.6241-4.2436 | 0.0232 |
| 0.03 | 0.0002 | 3.4941-3.7271 | 0.0053 | 35.7287-47.0914 | 0.0059 |
| 0.1 | 0.013 | 1.6786-1.9486 | 0.0012 | 37.5178-42.9054 | 0.0012 |
| 0.3 | 0.0215 | 1.1673-1.2809 | 0.009 | 34.5350-49.9702 | 0.0092 |
| 1 | 0.4306 | 0.9962-1.1483 | 0.0051 | 25.8500-33.8219 | 0.0048 |
| 5 | 0.0003 | 0.4797-0.6667 | 0.0014 | 18.9422-21.4334 | 0.0012 |
| 10 | 0.0263 | 0.3552-0.5607 | 0.0154 | 14.2808-22.4943 | 0.0129 |
| **LY500307** | | | | | |
| | **ERα** | | **ERβ** | | **ERα vs ERβ** |
| **Concentration (µmol/L)** | **p-value** | **95% CI** | **p-value** | **95% CI** | **p-value** |
| 0 | | 0.92-1.08 | | 0.83-1.17 | 1 |
| 0.01 | 0.0009 | 2.0095-2.2620 | 0.0037 | 73.7896-93.6231 | 0.0038 |
| 0.03 | 0.002 | 1.5881-1.7544 | 0.005 | 105.2260-107.7026 | 0.0051 |
| 0.1 | 0.0096 | 1.6203-1.9066 | 0.0102 | 63.9673-95.3686 | 0.0102 |
| 0.3 | 0.0029 | 1.8031-2.1447 | 0.0373 | 66.8043-80.2037 | 0.0372 |
| 1 | 0.0441 | 1.4546-2.0269 | 0.0019 | 67.8878-80.1651 | 0.0017 |
| 5 | 0.0221 | 1.4274-1.6103 | 0.0003 | 79.5217-79.9028 | 0.0014 |
| 10 | 0.046 | 1.1926-1.3828 | 0.0011 | 66.5517-77.7049 | 0.0011 |

**Supplemental Table 3A.** IC50 values, p-values , and 95% confidence intervals (CIs) were determined for cell viability in cytotoxicity assay with OSU-ERb-12 and LY500307. P-value (for IC50) for each cell line was determined by comparing with MCF10A.

| Cytotoxicity | | | | | | |
|---|---|---|---|---|---|---|
| | OSU-ERb-12 | | | LY500307 | | |
| Cell Line | IC50 (µmol/L) | p-value | 95% CI | IC50 (µmol/L) | p-value | 95% CI |
| MCF10A | 13.96 | | 7.186-19.64 | 30.53 | | 23.983-41.510 |
| MCF7 | 11.22 | 0.0750 | 9.578-13.10 | 3.48 | 0.0225 | 3.396-3.444 |
| MCF7-TamR | 9.27 | 0.0249 | 4.563-14.68 | 7.01 | 0.0286 | 6.678-7.215 |
| MCF7-FasR | 4.86 | 0.0087 | 2.643-8.555 | 1.67 | 0.0200 | 1.449-1.917 |
| MCF7-CDK6 OE | 10.75 | 0.0538 | 9.313-12.31 | 7.66 | 0.0288 | 6.766-8.409 |
| MCF7-CDK4/6iR | 7.18 | 0.0089 | 5.391-9.214 | 6.89 | 0.0337 | 6.714-7.066 |
| T47D | 10.43 | 0.0403 | 9.715-11.29 | 7.29 | 0.0297 | 7.260-7.560 |
| T47D-TamR | 9.40 | 0.0301 | 9.439-9.461 | 6.66 | 0.0258 | 5.281-7.925 |
| T47D-FasR | 9.48 | 0.0304 | 9.457-9.483 | 7.75 | 0.0307 | 7.693-7.973 |
| T47D-CDK4/6iR | 9.08 | 0.0245 | 8.878-9.042 | 7.47 | 0.0281 | 6.230-7.838 |

**Supplemental Table 3B.** P-values were determined for cell viability in cytotoxicity assay with OSU-ERb-12 and LY500307. P-value (for IC50) for each cell resistant cell line was determined by comparing with corresponding parental cell line (MCF7 or T47D).

| Cytotoxicity (parental vs drug resistant) | | |
|---|---|---|
| | OSU-ERb-12 | LY500307 |
| Cell Line | p-value | p-value |
| MCF7 | | |
| MCF7-TamR | 0.0118 | 0.0014 |
| MCF7-FasR | 0.0483 | 0.0043 |
| MCF7-CDK6 OE | 0.1302 | 0.0106 |
| MCF7-CDK4/6iR | 0.0031 | 0.0421 |
| T47D | | |
| T47D-TamR | 0.0415 | 0.3096 |
| T47D-FasR | 0.0478 | 0.0915 |
| T47D-CDK4/6iR | 0.0271 | 0.4181 |

**Supplemental Table 4.** IC50 values, p-values , and 95% confidence intervals (CIs) were determined for viability of T47D cells in cytotoxicity assay with OSU-ERb-12, LY500307 and combination of drugs as indicated. P-value (for IC50) for each combination was determined by comparing with OSU-ERb-12 or LY500307 alone (as indicated).

| T47D Cytotoxicity | | | |
|---|---|---|---|
| **Drug** | **IC50 (µmol/L)** | **p-value** | **95% CI** |
| OSU-ERb-12 | 14.10 | | 14.035-14.271 |
| OSU-ERb-12+ 0.5µmol/L 4OH-Tam | 1.00 | <0.0001 | 0.6828-1.53 |
| OSU-ERb-12+ 0.25µmol/L Fas | 4.47 | 0.0179 | 2.789-8.391 |
| OSU-ERb-12+ 1µmol/L Elacestrant | 6.18 | <0.0001 | 5.450-7.024 |
| OSU-ERb-12+ 0.75µmol/L MPP | 3.29 | 0.0038 | 2.646-4.170 |

| T47D Cytotoxicity | | | |
|---|---|---|---|
| **Drug** | **IC50 (µmol/L)** | **p-value** | **95% CI** |
| OSU-ERb-12 | 10.41 | | 9.715-11.29 |
| OSU-ERb-12+0.75µmol/L MPP | 1.22 | 0.0005 | 0.8112-1.823 |
| LY500307 | 7.29 | | 7.131-7.708 |
| LY500307+0.75µmol/L MPP | 4.48 | 0.001 | 4.346-4.598 |

**Supplemental Table 5.** Cell viability, Bliss prediction, synergy ratio, synergy 95% CI, and p-values were determined at each drug combination (in T47D cells) as indicated.

| Bliss Independence Model | | | | | |
|---|---|---|---|---|---|
| **Combination** | **Combination Viability** | **Bliss prediction** | **Synergy Ratio** | **Synergy 95% CI** | **p-value** |
| 0.125 µmol/L OSU-ERb-12 + 0.025 µmol/L Tam | 0.83 | 1.59 | 1.9 | 1.44-2.51 | 0.001 |
| 0.250 µmol/L OSU-ERb-12 +0.0 50 µmol/L Tam | 0.83 | 1.54 | 1.84 | 1.42-2.39 | 0.001 |
| 0.500 µmol/L OSU-ERb-12 + 0.100 µmol/L Tam | 0.86 | 1.8 | 2.1 | 1.66-2.65 | <0.001 |
| 1.25 µmol/L OSU-ERb-12 + 0.250 µmol/L Tam | 0.81 | 1.75 | 2.16 | 1.52-3.05 | 0.002 |
| 2.5 µmol/L OSU-ERb-12 + 0.500 µmol/L Tam | 0.82 | 1.97 | 2.41 | 1.85-3.13 | <0.001 |
| 5 µmol/L OSU-ERb-12 + 1 µmol/L Tam | 0.54 | 1.27 | 2.35 | 1.83-3.02 | <0.001 |

**Supplemental Table 6.** IC50 values, p-values , and 95% confidence intervals (CIs) were determined for cell viability in cytotoxicity assay (in T47D cells) with OSU-ERb-12 alone, MCSR-18-006 alone and combination as indicated. P-value (for IC50) for OSU-ERb-12 alone/combination were determined by comparing with MCSR-18-006 alone or respective combination treatment.

| T47D Cytotoxicity | | | |
|---|---|---|---|
| Drug | IC50 (μmol/L) | p-value | 95% CI |
| MCSR-18-006 | 33.68 | | 30.82-36.64 |
| OSU-ERb-12 | 10.41 | 0.0026 | 9.715-11.29 |

| T47D Cytotoxicity | | | |
|---|---|---|---|
| Drug | IC50 (μmol/L) | p-value | 95% CI |
| MCSR-18-006+0.5μmol/L 4-OH-Tam | 39.23 | | 36.56-41.01 |
| OSU-ERb-12+0.5μmol/L 4-OH-Tam | 1.02 | <0.0001 | 0.6044-1.552 |

**Supplemental Table 7.** p-values and 95% confidence intervals (CIs) were determined for proliferation of MCF7 and T47D cells. P-value was determined by comparing with the percentage of cell proliferation in each drug treatment with that of the corresponding control (in each cell line).

| Proliferation | | | | | |
|---|---|---|---|---|---|
| MCF7 | | | T47D | | |
| Treatment | p-value | 95% CI | Treatment | p-value | 95% CI |
| Control | | 50.125-53.475 | Control | | 46.355-48.978 |
| Fas 0.5μmol/L | 0.002 | 18.996-23.504 | Fas 0.5μmol/L | 0.0033 | 26.163-28.237 |
| OSU-ERb-12 0.5μmol/L | 0.0096 | 36.073-51.861 | OSU-ERb-12 0.5 μmol/L | 0.0087 | 54.730-61.137 |
| OSU-ERb-12 10μmol/L | 0.0163 | 23.900-26.167 | OSU-ERb-12 10 μmol/L | 0.0074 | 12.722-20.678 |
| Control | | 24.232-28.035 | Control | | 46.355-48.978 |
| Fas 0.5 μmol/L | 0.001 | 12.180-13.687 | Fas 0.5μmol/L | 0.0189 | 26.796-33.871 |
| LY500307 0.5 μmol/L | 0.2491 | 39.082-41.918 | LY500307 0.5μmol/L | 0.02 | 57.681-62.986 |
| LY500307- 3 μmol/L | 0.0028 | 4.483-10.250 | LY500307- 7μmol/L | 0.0154 | 30.541-35.192 |

**Supplemental Table 8.** p-values and 95% confidence intervals (CIs) were determined for each phase of cell cycle in OSU-ERb-12- and LY500307-treated MCF7 and T47D cells. P-value for each phase for each treatment was determined by comparing with the percentage of cells present in the corresponding control (DMSO treated) cells.

| Cell Cycle | | | | | | | | |
|---|---|---|---|---|---|---|---|---|
| **MCF7** | | | | | | | | |
| | **Sub G0** | | **G0/G1** | | **S** | | **G2/M** | |
| **Treatment** | **p-value** | **95% CI** | **p-value** | **95% CI** | **p-value** | **95% CI** | **p-value** | **95% CI** |
| Control | | - | | 66.396-67.338 | | 13.971-15.896 | | 15.313-17.354 |
| OSU-ERb-12 0.5µmol/L | - | - | 0.0200 | 55.239-61.028 | 0.0059 | 21.098-21.569 | 0.0890 | 16.706-21.561 |
| OSU-ERb-12 11µmol/L | 1.0000 | 0.087-0.313 | 0.0038 | 75.655-78.145 | 0.0347 | 9.219-11.581 | 0.0020 | 10.239-11.694 |
| LY500307 0.5µmol/L | 0.2254 | 0.187-0.413 | 0.0191 | 49.863-57.937 | 0.0493 | 19.719-24.414 | 0.0097 | 20.821-21.512 |
| LY500307 3µmol/L | 0.1221 | 0.338-1.195 | 0.4598 | 66.517-68.216 | 0.2556 | 14.843-16.090 | 0.2965 | 15.172-15.695 |
| **T47D** | | | | | | | | |
| | **Sub G0** | | **G0/G1** | | **S** | | **G2/M** | |
| **Treatment** | **p-value** | **95% CI** | **p-value** | **95% CI** | **p-value** | **95% CI** | **p-value** | **95% CI** |
| Control | | 0.305-1.162 | | 81.475-82.858 | | 6.794-7.139 | | 7.7664-9.803 |
| OSU-ERb-12 0.5µmol/L | 0.1942 | 1.084-4.382 | 0.0036 | 74.582-76.685 | 0.0015 | 11.931-12.402 | 1.0000 | 7.719-9.748 |
| OSU-ERb-12 10µmol/L | 0.2799 | 0.317-3.017 | 0.6010 | 84.704-82.096 | 0.0130 | 8.903-9.630 | 0.0749 | 6.199-7.867 |
| LY500307 0.5µmol/L | 0.2107 | 0.956-4.244 | 0.0018 | 73.166-75.834 | 0.0004 | 13.168-13.299 | 0.8758 | 7.779-9.954 |
| LY500307 7µmol/L | 0.0068 | 5.781-6.819 | 0.0079 | 42.248-53.019 | 0.0060 | 17.673-21.860 | 0.0135 | 15.508-17.092 |

**Supplemental Table 9.** p-values and 95% confidence intervals (CIs) were determined for induction of apoptosis in OSU-ERb-12- and LY500307-treated MCF7 and T47D cells. P-value (for apoptosis induction) for each treatment was determined by comparing with the percentage of apoptotic cells present in the corresponding control (DMSO treated) cells.

| Apoptosis | | | | | |
|---|---|---|---|---|---|
| **MCF7** | | | **T47D** | | |
| **Treatment** | **p-value** | **95% CI** | **Treatment** | **p-value** | **95% CI** |
| Control | | 4.56-6.64 | Control | | 5.16-7.84 |
| OSU-ERb-12 (0.5µmol/L) | 0.4 | 6.03-7.97 | OSU-ERb-12 (0.5µmol/L | 0.3 | 4.77-7.29 |
| OSU-ERb-12 (10µmol/L) | 0.04 | 7.45-9.95 | OSU-ERb-12 (10µmol/L) | 0.0226 | 12-13.9 |
| Control | | 4.56-6.64 | Control | | 2.23-4.17 |
| LY500307 (0.5µmol/L) | 0.03 | 6.03-7.97 | LY500307 (0.5µmol/L) | 0.003 | 8.88-11.4 |
| LY500307 (3µmol/L) | 0.015 | 7.45-9.95 | LY500307 (7µmol/L) | 0.0005 | 9.26-13 |

**Supplemental Table 10.** p-values and 95% confidence intervals (CIs) were determined for assessing the colony forming ability in OSU-ERb-12- and LY500307-treated MCF7 and T47D cells. P-value (for percentage of colony formation) for each treatment was determined by comparing with the percentage of that in the corresponding control (DMSO treated) cells.

| Colony Formation Assay | | | | |
|---|---|---|---|---|
| | MCF7 | | T47D | |
| Treatment | p-value | 95% CI | p-value | 95% CI |
| Control | | 560.6272-622.0394 | | 409.86-532.14 |
| OSU-ERb-12 (3µmol/L) | 0.050162 | 502.12-515.88 | 0.180841 | 533.9786-569.3546 |
| OSU-ERb-12 (5µmol/L) | 0.001899 | 315.0875-350.2457 | 0.011064 | 107.4305-227.2361 |
| LY500307 (3µmol/L) | 0.003087 | 96.5738-152.7594 | 0.014908 | 316.7503-437.9163 |
| LY500307 (5µmol/L) | 0.000701 | 0 | 0.005 | 21.04-24.93 |

**Supplemental Table 11.** p-values and 95% confidence intervals (CIs) were determined for assessing the cell migration ability in OSU-ERb-12- and LY500307-treated MCF7 cells. P-value (for percentage of cell migration) for each treatment was determined by comparing with the percentage of that in the corresponding control (DMSO treated) cells.

| Cell Migration Assay | | |
|---|---|---|
| | 24 Hours | |
| Treatment | p-value | 95% CI |
| Control | | 91.76-108.24 |
| OSU-ERb-12 (5µmol/L) | 0.0004368 | 61.0140-69.5791 |
| OSU-ERb-12 (10µmol/L) | 0.0026058 | 52.0192-62.1763 |
| LY500307 (5µmol/L) | 8.438E-06 | 23.4626-36.1402 |
| LY500307 (10µmol/L) | 2.756E-05 | 5.1938-11.0745 |

**Supplemental Table 12:** Patient demographics.

| | Overall (n=37) |
|---|---|
| **Age at diagnosis** | |
| **Median [min, max]** | 56 [35, 169] |
| **Race** | |
| **Asian** | 1 (3%) |
| **Black or African American** | 1 (3%) |
| **White** | 35 (95%) |
| **Ethnicity** | |
| **NOT Hispanic or Latino** | 37 (100%) |
| **BMI** | |
| **Median [min, max]** | 28 [19, 47] |
| **Stage (pathologic)** | |
| **I** | 5 (14%) |
| **II** | 8 (22%) |
| **III** | 15 (41%) |
| **IV** | 7 (19%) |
| **Unknown** | 2 (5%) |
| **Grade** | |
| **High grade (3)** | 7 (19%) |
| **Intermediate grade (2)** | 23 (62%) |
| **Low grade (1)** | 4 (11%) |
| **Unknown** | 3 (8%) |
| **ER status** | |
| **Positive** | 37 (100%) |
| **PR status** | |
| **Negative** | 4 (11%) |
| **Positive** | 33 (89%) |
| **HER2 status** | |
| **Equivocal** | 2 (5%) |
| **Negative** | 34 (92%) |
| **Positive** | 1 (3%) |
| **Chemotherapy** | |
| **No** | 8 (22%) |
| **Yes** | 29 (78%) |
| **Hormone therapy** | |
| **No** | 1 (3%) |
| **Yes** | 36 (97%) |
| **Targeted therapy** | |
| **No** | 20 (54%) |
| **Yes** | 17 (46%) |

### Figure S1

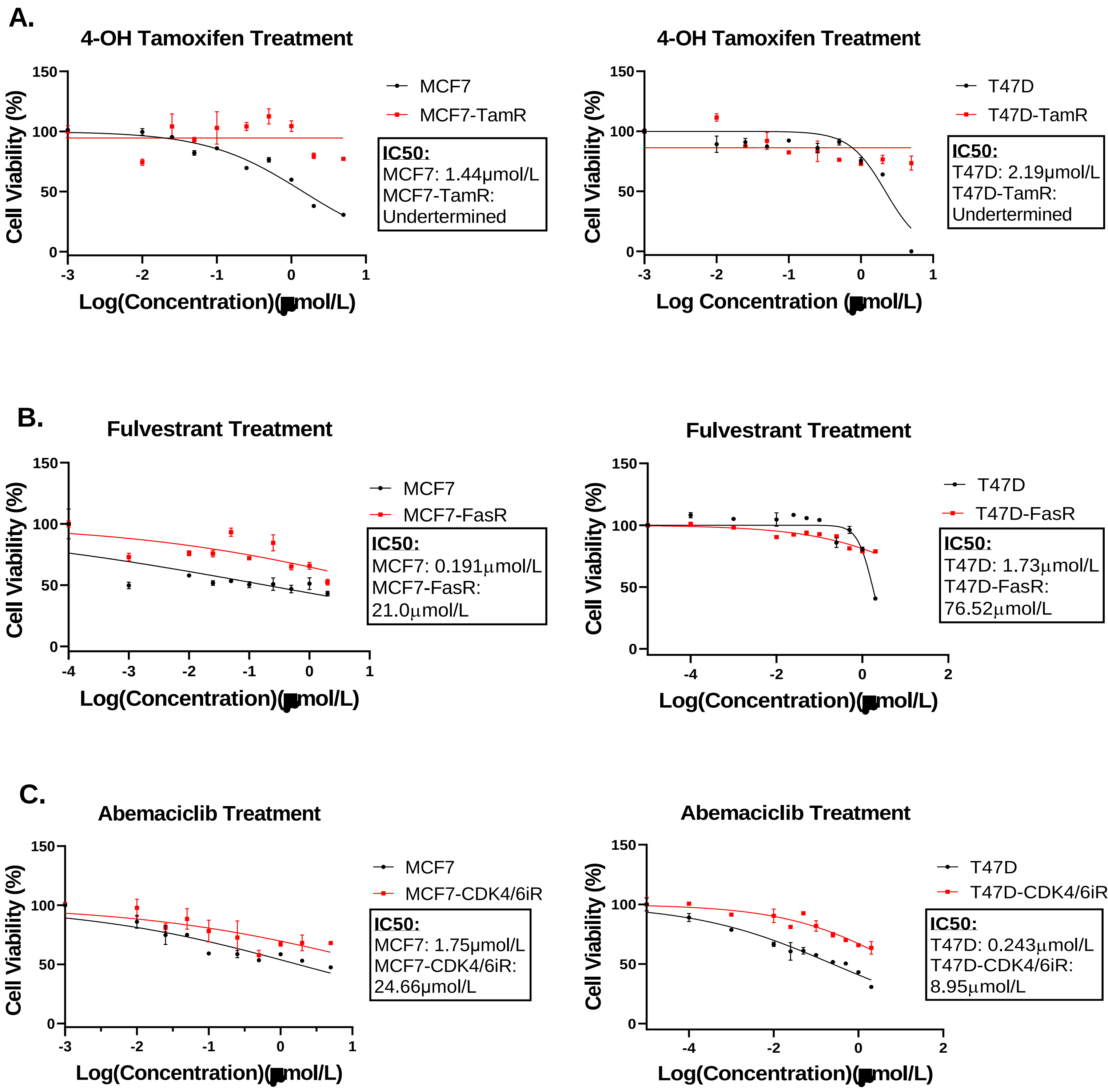
A.
4-OH Tamoxifen Treatment
Cell Viability (%)
Log(Concentration)(µmol/L)
MCF7
MCF7-TamR
IC50:
MCF7: 1.44μmol/L
MCF7-TamR:
Undertermined
4-OH Tamoxifen Treatment
Cell Viability (%)
Log Concentration (µmol/L)
T47D
T47D-TamR
IC50:
T47D: 2.19μmol/L
T47D-TamR:
Undertermined
B.
Fulvestrant Treatment
Cell Viability (%)
Log(Concentration)(µmol/L)
MCF7
MCF7-FasR
IC50:
MCF7: 0.191µmol/L
MCF7-FasR:
21.0µmol/L
Fulvestrant Treatment
Cell Viability (%)
Log(Concentration)(µmol/L)
T47D
T47D-FasR
IC50:
T47D: 1.73µmol/L
T47D-FasR:
76.52µmol/L
C.
Abemaciclib Treatment
Cell Viability (%)
Log(Concentration)(µmol/L)
MCF7
MCF7-CDK4/6iR
IC50:
MCF7: 1.75μmol/L
MCF7-CDK4/6iR:
24.66μmol/L
Abemaciclib Treatment
Cell Viability (%)
Log(Concentration)(µmol/L)
T47D
T47D-CDK4/6iR
IC50:
T47D: 0.243µmol/L
T47D-CDK4/6iR:
8.95µmol/L

### Figure S2

**OSU-ERb-12**

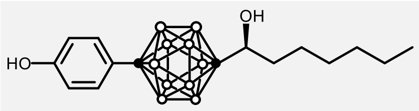

(*R*)-1-[1-(4-Hydroxyphenyl)-1,12-dicarba-*closo*-dodecaborane-12-yl]heptan-1-ol.

**MCSR-18-006**

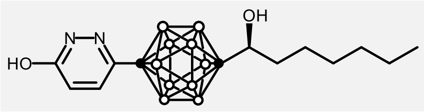

Black dots are carbon, white dots represent B-H

**LY500307**

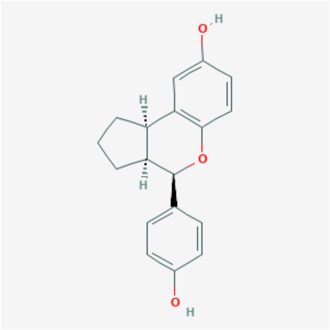

**4-OH-Tamoxifen**

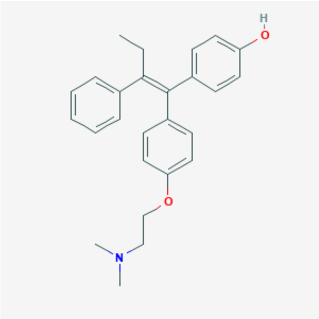

**Elacestrant**

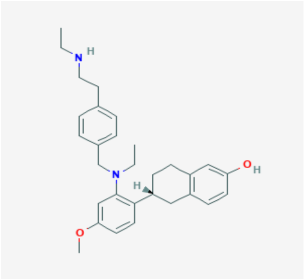

**Fulvestrant**

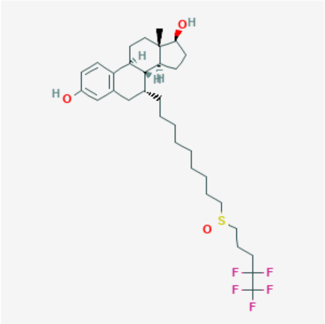

**DPN (Diarylpropionitrile)**

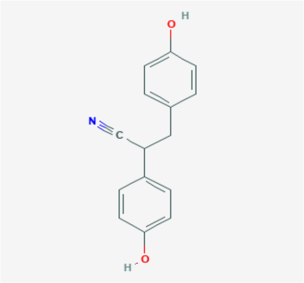

**AC186**

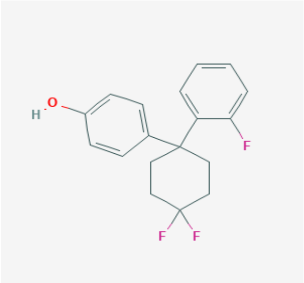

**WAY 200070**

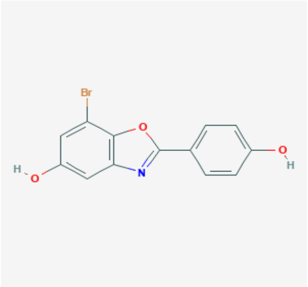

**MPP (Methyl-piperidino-pyrazole)**

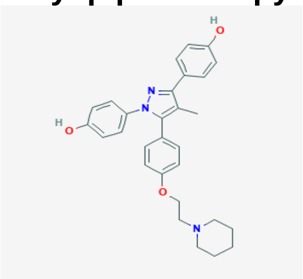

**Figure S3**

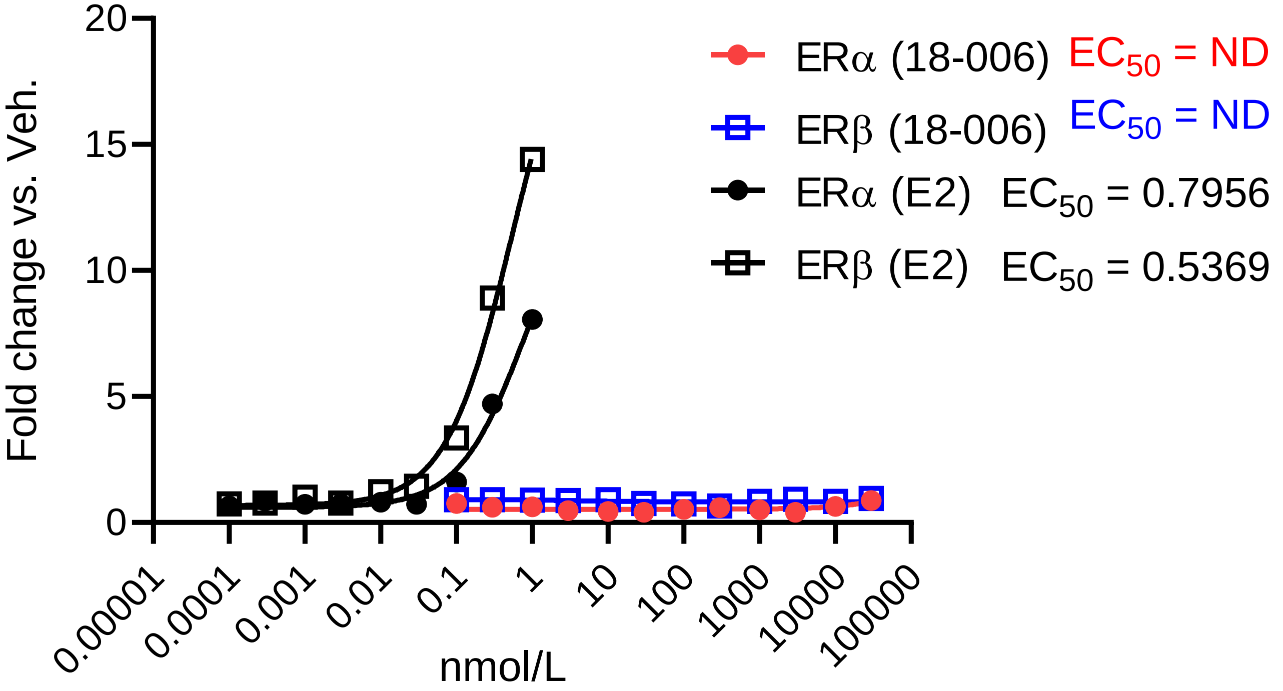

Fold change vs. Veh.
20
15
10
5
0
0.00001
0.0001
0.001
0.01
0.1
1
10
100
1000
10000
100000
nmol/L
ERα (18-006) $EC_{50}$ = ND
ERβ (18-006) $EC_{50}$ = ND
ERα (E2) $EC_{50}$ = 0.7956
ERβ (E2) $EC_{50}$ = 0.5369

#### Figure S4

### Radio-ligand Displacement Assay

**DES (+) control**

#### Reference Compound

| Assay Name | Reference Compound | $IC_{50}$ |
|---|---|---|
| Estrogen ERα | Diethylstilbestrol | 1.32 nmol/L |
| Estrogen ERβ | Diethylstilbestrol | 1.06 nmol/L |

ERα: IC50 = 1.32 nmol/L, Ki = 0.38 nmol/L
ERβ: IC50 = 1.06 nmol/L, Ki = 0.22 nmol/L

### Radio-ligand Displacement Assay

**MCSR-18-006: Test Compound**

#### Test Compound
#### Experimental Results

| Assay Name | Batch * | Spec. | Rep. | Conc. | % Inh. | $IC_{50}$* | $K_i$ |
|---|---|---|---|---|---|---|---|
| Estrogen ERα | 461679 | hum | 2 | 100 µmol/L | 24 | >100 µmol/L | |
| | | hum | 2 | 10 µmol/L | 6 | | |
| | | hum | 2 | 1 µmol/L | 3 | | |
| | | hum | 2 | 0.1 µmol/L | 8 | | |
| | | hum | 2 | 10 nmol/L | -6 | | |
| | | hum | 2 | 1 nmol/L | 3 | | |
| | | hum | 2 | 0.1 nmol/L | 1 | | |
| | | hum | 2 | 10 pmol/L | 4 | | |
| Estrogen ERβ | 461680 | hum | 2 | 100 µmol/L | 39 | >100 µmol/L | |
| | | hum | 2 | 10 µmol/L | 8 | | |
| | | hum | 2 | 1 µmol/L | -2 | | |
| | | hum | 2 | 0.1 µmol/L | 0 | | |
| | | hum | 2 | 10 nmol/L | -7 | | |
| | | hum | 2 | 1 nmol/L | 5 | | |
| | | hum | 2 | 0.1 nmol/L | 4 | | |
| | | hum | 2 | 10 pmol/L | -3 | | |

#### Figure S5

**A.**

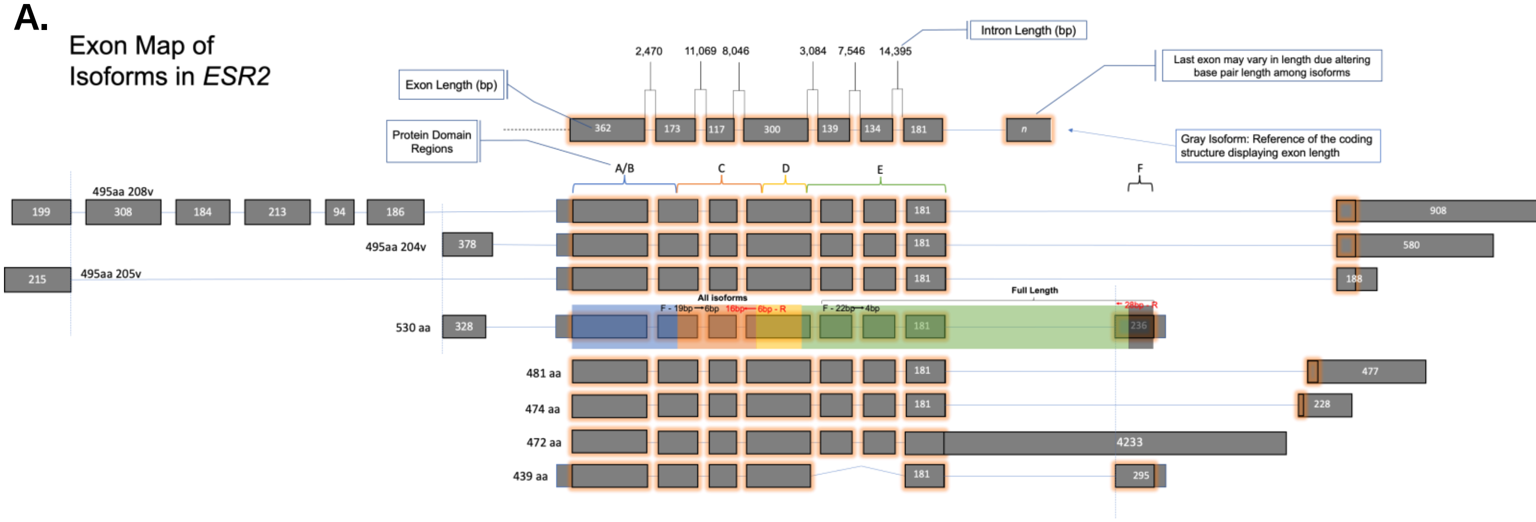
Exon Map of Isoforms in ESR2
Intron Length (bp)
2,470
11,069
8,046
3,084
7,546
14,395
Exon Length (bp)
Last exon may vary in length due altering base pair length among isoforms
Protein Domain Regions
Gray Isoform: Reference of the coding structure displaying exon length
362
173
117
300
139
134
181
n
A/B
C
D
E
F
495aa 208v
199
308
184
213
94
186
908
495aa 204v
378
580
215
495aa 205v
188
Full Length
All isoforms
F - 19bp → 6bp
16bp ← 6bp - R
F - 22bp → 4bp
← 28bp - R
530 aa
328
236
481 aa
477
474 aa
228
472 aa
4233
439 aa
295

**B.**

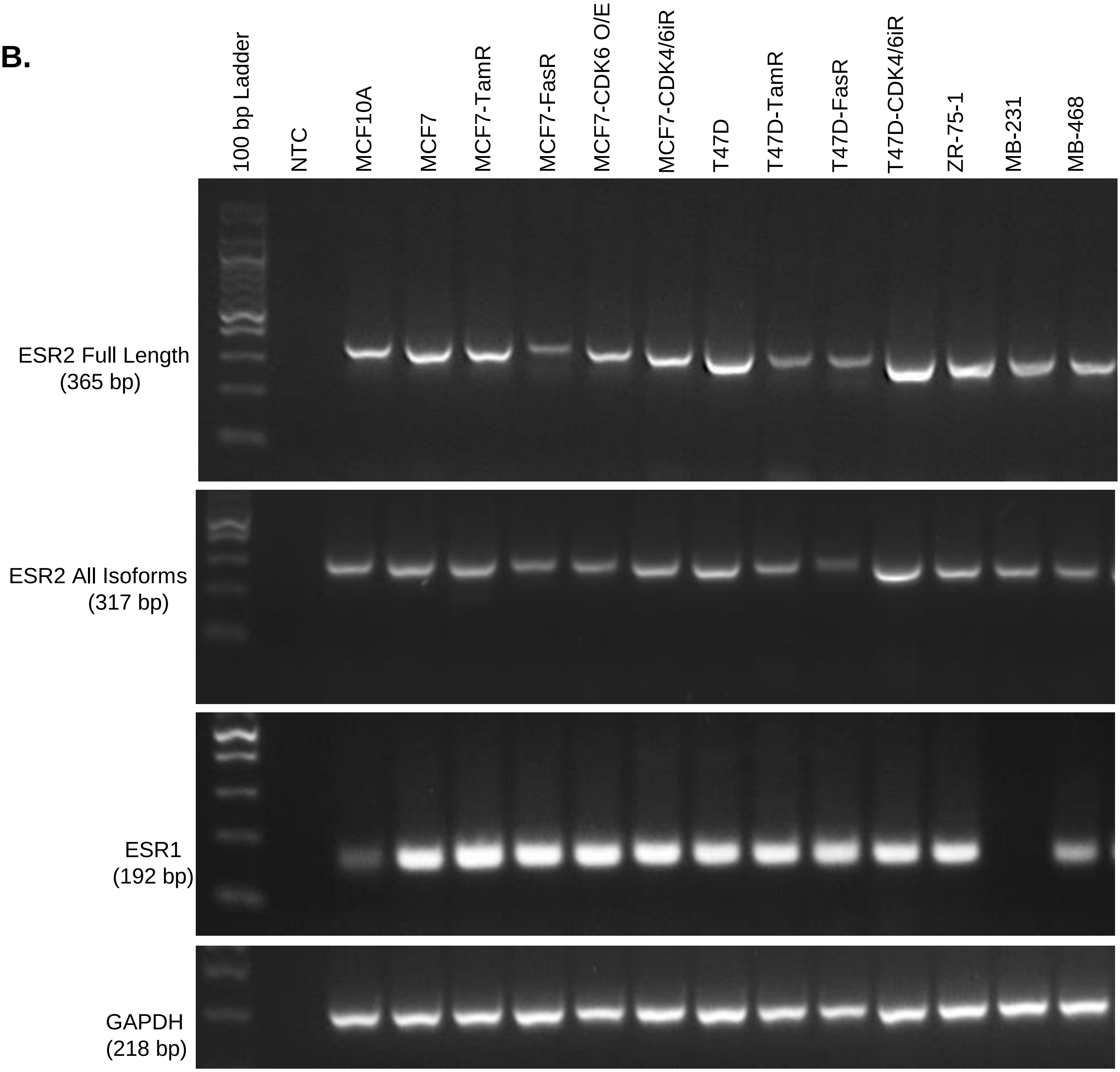
100 bp Ladder
NTC
MCF10A
MCF7
MCF7-TamR
MCF7-FasR
MCF7-CDK6 O/E
MCF7-CDK4/6iR
T47D
T47D-TamR
T47D-FasR
T47D-CDK4/6iR
ZR-75-1
MB-231
MB-468
ESR2 Full Length (365 bp)
ESR2 All Isoforms (317 bp)
ESR1 (192 bp)
GAPDH (218 bp)

**Figure S6**

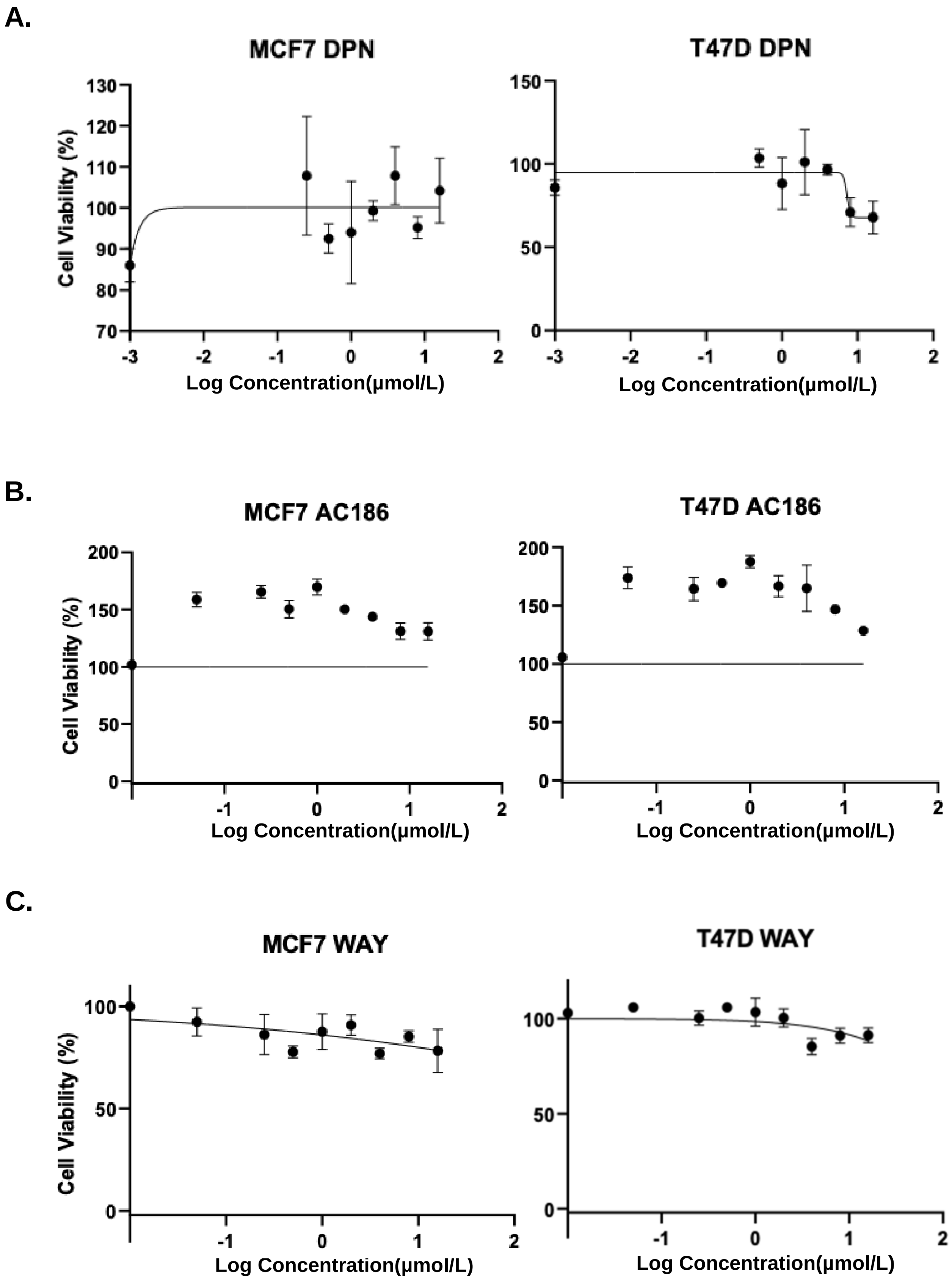
A.
MCF7 DPN
T47D DPN
B.
MCF7 AC186
T47D AC186
C.
MCF7 WAY
T47D WAY
Cell Viability (%)
Log Concentration(μmol/L)

### Figure S7

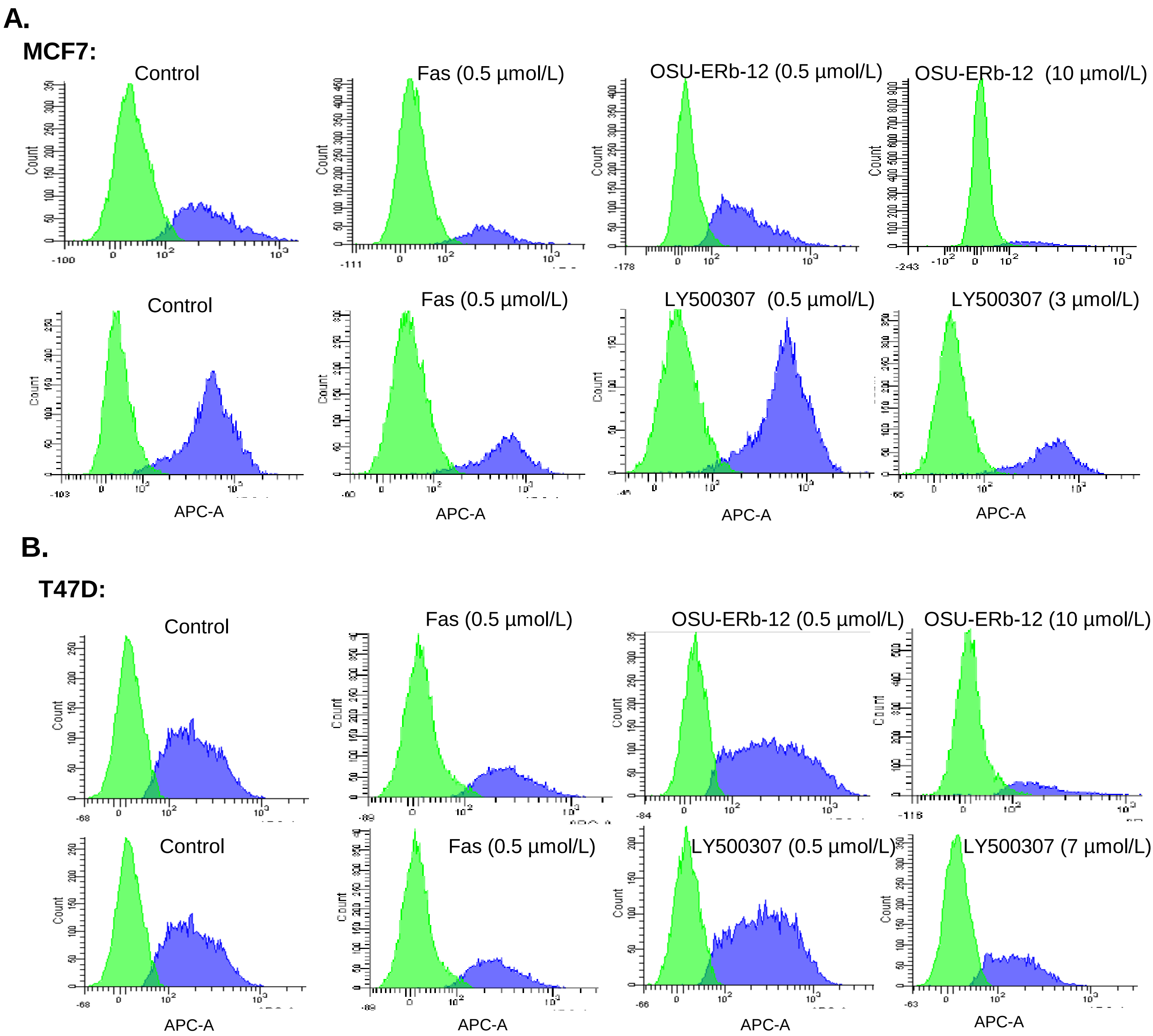
A.
MCF7:
Control
Fas (0.5 µmol/L)
OSU-ERb-12 (0.5 µmol/L)
OSU-ERb-12 (10 µmol/L)
Control
Fas (0.5 µmol/L)
LY500307 (0.5 µmol/L)
LY500307 (3 µmol/L)
Count
APC-A
B.
T47D:
Control
Fas (0.5 µmol/L)
OSU-ERb-12 (0.5 µmol/L)
OSU-ERb-12 (10 µmol/L)
Control
Fas (0.5 µmol/L)
LY500307 (0.5 µmol/L)
LY500307 (7 µmol/L)
Count
APC-A

### Figure S8

#### Cell Cycle Analysis

**A.**

**MCF7:**

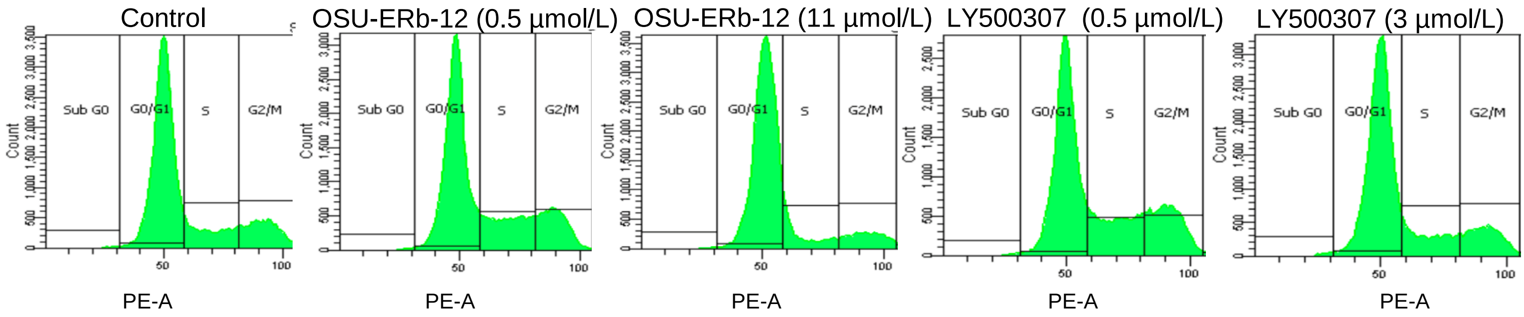

**B.**

**T47D:**

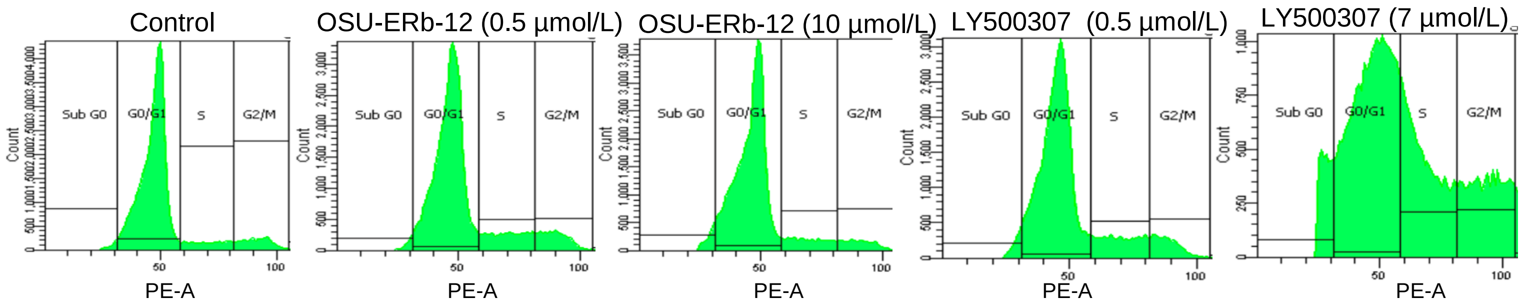

### Figure S9

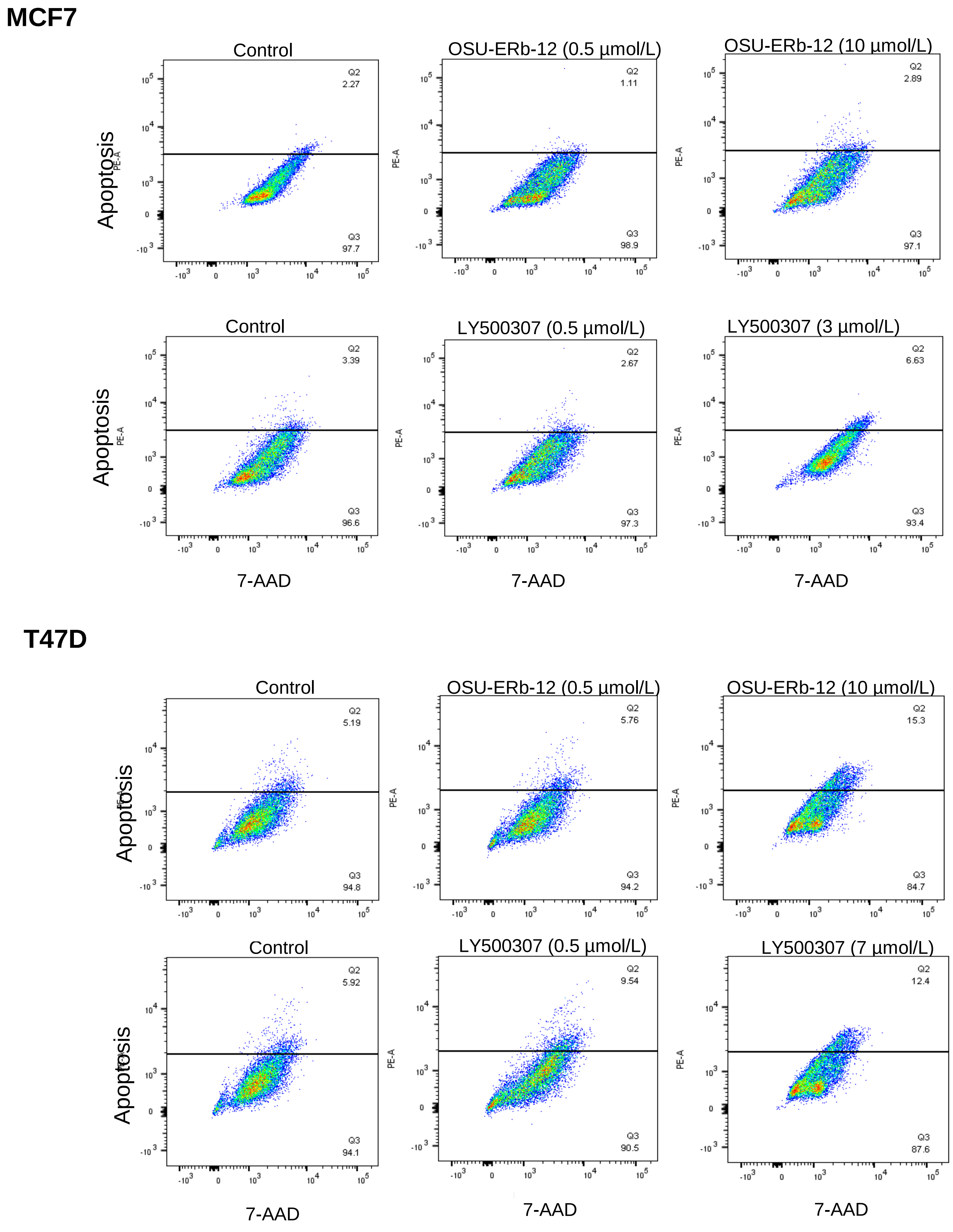
MCF7
Control
Q2
2.27
Q3
97.7
OSU-ERb-12 (0.5 μmol/L)
Q2
1.11
Q3
98.9
OSU-ERb-12 (10 μmol/L)
Q2
2.89
Q3
97.1
Control
Q2
3.39
Q3
96.6
LY500307 (0.5 μmol/L)
Q2
2.67
Q3
97.3
LY500307 (3 μmol/L)
Q2
6.63
Q3
93.4
Apoptosis
PE-A
7-AAD
T47D
Control
Q2
5.19
Q3
94.8
OSU-ERb-12 (0.5 μmol/L)
Q2
5.76
Q3
94.2
OSU-ERb-12 (10 μmol/L)
Q2
15.3
Q3
84.7
Control
Q2
5.92
Q3
94.1
LY500307 (0.5 μmol/L)
Q2
9.54
Q3
90.5
LY500307 (7 μmol/L)
Q2
12.4
Q3
87.6
Apoptosis
PE-A
7-AAD
